# Neuroimaging within the Dominantly Inherited Alzheimer’s Network (DIAN): PET and MRI

**DOI:** 10.1101/2022.03.25.485799

**Authors:** Nicole S. McKay, Brian A. Gordon, Russ C. Hornbeck, Clifford R. Jack, Robert Koeppe, Shaney Flores, Sarah Keefe, Diana A. Hobbs, Nelly Joseph-Mathurin, Qing Wang, Farzaneh Rahmani, Charles D. Chen, Austin McCullough, Deborah Koudelis, Jasmin Chua, Beau M. Ances, Peter R. Millar, Mike Nickels, Richard J. Perrin, Ricardo F. Allegri, Sarah B. Berman, William S. Brooks, David M. Cash, Jasmeer P. Chhatwal, Martin R. Farlow, Nick C. Fox, Michael Fulham, Berhadino Ghetti, Neill Graff-Radford, Takeshi Ikeuchi, Gregg Day, William Klunk, Johannes Levin, Jae-Hong Lee, Ralph Martins, Colin L. Masters, Jonathan McConathy, Hiroshi Mori, James M. Noble, Christopher Rowe, Stephen Salloway, Raquel Sanchez-Valle, Peter R. Schofield, Hiroyuki Shimada, Mikio Shoji, Yi Su, Kazushi Suzuki, Jonathan Vöglein, Igor Yakushev, Laura Swisher, Carlos Cruchaga, Jason Hassenstab, Celeste Karch, Eric McDade, Chengjie Xiong, John C. Morris, Randall J. Bateman, Tammie L.S. Benzinger, Dominantly Inherited Alzheimer Network

**Author notes:** Corresponding author: Tammie L.S. Benzinger Mallinckrodt Institute of Radiology, Washington University in Saint Louis, Saint Louis, Missouri, USA.

## Abstract

The Dominantly Inherited Alzheimer Network (DIAN) Observational Study is an international collaboration studying autosomal dominant Alzheimer disease (ADAD). This rare form of Alzheimer disease (AD) is caused by mutations in the presenilin 1 *(PSEN1)*, presenilin 2 *(PSEN2)*, or amyloid precursor protein (*APP*) genes. As individuals from these families have a 50% chance of inheriting the familial mutation, this provides researchers with a well-matched cohort of carriers vs non-carriers for case-control studies. An important trait of ADAD is that the age at symptom onset is highly predictable and consistent for each specific mutation, allowing researchers to estimate an individual’s point in their disease time course prior to symptom onset. Although ADAD represents only a small proportion (approximately 0.1%) of all AD cases, studying this form of AD allows researchers to investigate preclinical AD and the progression of changes that occur within the brain prior to AD symptom onset. Furthermore, the young age at symptom onset (typically 30-60 years) means age-related comorbidities are much less prevalent than in sporadic AD, thereby allowing AD pathophysiology to be studied independent of these confounds. A major goal of the DIAN Observational Study is to create a global resource for AD researchers. To that end, the current manuscript provides an overview of the DIAN magnetic resonance imaging (MRI) and positron emission tomography (PET) protocols and highlights the key imaging results of this study to date.

## Introduction

Understanding the brain in health and in neurodegenerative disease has broad implications for public health, and researchers are increasingly forming collaborations across institutions to study these disorders. As populations continue to age, Alzheimer disease (AD) is a pressing research priority increasingly recognized as a global problem that requires an international response and research collaborations ^1^. Therefore, it is unsurprising that some of the earliest and most well-established collaborative efforts have focused on this disease. The Alzheimer Disease Neuroimaging Initiative (ADNI) ^2^ and the National Alzheimer’s Coordinating Center (NACC) ^3^ are comprised of large networks of researchers who are working together to study the progression of, and potential therapeutic treatments for, this complex and devastating disease. More specialized consortia also exist for researchers studying specific subtypes of AD including the Longitudinal Early-Onset Alzheimer’s Disease Study (LEADS) ^4^, or populations who are highly susceptible to this disease, such as the Alzheimer Biomarker Consortium-Down Syndrome (ABC-DS) ^5^. One such group, the Dominantly Inherited Alzheimer Network (DIAN) ^6, 7^, brings together researchers at 22 institutes across Asia, Australasia, Europe, and the Americas, to study ADAD.

ADAD is a rare form of AD, accounting for about 0.1% of all AD cases ^8, 9^, that occurs as a result of mutations or duplications in three genes; amyloid precursor protein *(APP),* presenilin 1 *(PSEN1),* and presenilin 2 *(PSEN2)* ^10–13^. In total more than 230 different mutations have been identified within the genes coding for these proteins ^14^. Mechanistically these mutations cause the increased aggregation of beta-amyloid (Aβ) in the brain parenchyma by either increasing the overall production of Aβ or by relative increases in aggregation-prone Aβ subtypes ^15, 16^ that then triggers a cascade of events that lead individuals on a pathway to developing symptomatic AD ^17, 18^. Individuals with ADAD typically show first symptoms at a young age. First symptoms are typically reported when an individual is 30-50 years old, however, this age varies by mutation and may occur earlier or later ^6^. In contrast, sporadic AD develops via a complex interaction of genetic, lifestyle, and environmental influences. These multifaceted causes of sporadic AD impact both the production and clearance of Aβ ^19^ and typically affect adults 65 years and older, although early onset forms of sporadic AD do exist.

As with sporadic forms of the disease, ADAD mutation-carriers accumulate the cerebral Aβ plaques and tau neurofibrillary tangles that are the hallmarks of AD ^20–22^, although burdens of neuropathologic lesions are typically greater in ADAD at expiration ^23, 24^. Given that individuals with ADAD are often much younger, they harbor fewer common age-related pathologies. In post-mortem examinations, sporadic AD cases commonly demonstrate vascular or other co-pathology of lacunes, atherosclerosis of the Circle of Willis, and parenchymal arteriosclerosis, amongst many others ^25^. Such age-related pathologies are rare in ADAD, although Lewy Body and Transactive DNA-binding protein 43 (TDP-43) proteinopathies do co-occur with AD pathology in ADAD ^26^. ADAD also has a higher prevalence of cerebral amyloid angiopathy ^24^, particularly in the Dutch and Flemish variants of the mutations ^20–22^. Importantly, the relatively low occurrence of AD-independent age-related neuropathologies allows for a greater understanding of how AD processes impact the brain structurally and functionally, with a reduction in the confounding effect of age.

Beyond the important role studying ADAD has had in shaping our understanding of crucial elements of AD pathophysiology, there are several remarkable advantages to the longitudinal study of ADAD. For example, the lack of age-related co-morbidities in ADAD allow AD pathology to be more directly linked to biomarker and clinical changes without the typical confounds of vascular or other pathologies reported in sporadic AD populations.

There is also certainty about the causative pathology of ADAD prior to autopsy, which allows for more secure inferences to be made in research focusing on understanding ADAD *in vivo*. Furthermore, ADAD has a relatively predictable age at symptom onset allowing for presymptomatic changes to be “staged” relative to the expected age at onset without needing to wait for an individual to become symptomatic. The 50% chance of becoming symptomatic combined with the predictable age at onset allows research studies to deliver adequate power with fewer enrolled participants. Finally, a major advantage of studying ADAD is that it allows us the ability to relate changes very closely to known drivers of the disease process such as specific mutations. In contrast, this is not possible in sporadic AD where one does not know how much APOE, head injury, cholesterol, or other lifestyle factors, may be driving disease onset or modifying the expressed phenotype.

Using positron emission tomography (PET), it has been reported that extensive Aβ plaque deposits accumulate in individuals with ADAD ^27, 28^. The topography of these accumulated plaques measured *in vivo* correspond strongly to patterns of Aβ reported in post-mortem studies ^29–31^. Across the disease course of AD, Aβ has been proposed to accumulate in a systematic pattern ^8, 32^. Studies comparing ADAD to sporadic AD indicate that patterns of Aβ deposition and spread may be topographically similar, although ADAD has noticeably elevated uptake in the basal ganglia, as well as greater general levels of tracer uptake ^27, 32–35^.

These differences in Aβ spatial patterns may reflect the mutation-specific changes in Aβ conformation present in these ADAD variants ^32, 36^, although basal ganglia deposition is also a feature found in Down Syndrome ^37^. The relationship between Aβ and the development of AD is complex, with recent work highlighting that many cognitively unimpaired older adults accumulate Aβ, and that this seems to moderate the relationship between brain structure and cognition ^38^. This complex relationship, and the clear evidence that Aβ accumulation likely starts a cascade of AD-specific pathology within the brain, has led many to propose Aβ as a potential therapeutic target of AD ^39–43^.

Similarly, the cerebral accumulation of tau-containing histopathological lesions are characteristic of AD ^44, 45^, and observable *in vivo* using tau-PET tracers ^46–48^. These lesions - neurofibrillary tangles (NFTs), neuritic plaques, and neuropil threads, together - are reflected in the intensity of the tau-PET signal. Spatial pattern analyses have shown that ADAD individuals have higher levels of tau-PET uptake in the precuneus and neocortex, compared to individuals with sporadic AD ^49, 50^. This increased tau pathology in ADAD is supported by post-mortem studies, which also find an abundance of tau pathology in these individuals ^23, 24^. Tau is also a candidate therapeutic target of AD ^51–53^, given that it is proposed to accumulate immediately prior to symptom onset ^8^, and is found to be highly predictive of cognitive decline ^52, 54, 55^ and structural atrophy ^56^.

In addition to abnormal Aβ and tau protein accumulation, neuroimaging provides the ability to measure non-specific neurodegenerative markers of AD such as hypometabolism and structural atrophy. [^18^F]-fluorodeoxyglucose (FDG) is a PET radiotracer sensitive to fluctuations in glucose metabolism, a measure of brain function ^57^. Reductions in glucose metabolism have been noted in individuals beginning up to 10 years before AD symptom onset ^8, 27, 58–60^, making hypometabolism an important biomarker for understanding the preclinical changes to brain function that happen in this disease. Previous studies using FDG-PET have reported global hypometabolism in ADAD individuals prior to cognitive decline, with the largest reductions in glucose metabolism in the temporoparietal, posterior cingulate, and medial temporal cortices ^61–64^. The degree of metabolic change observed in these regions has been shown to increase as individuals approach their EYO, while the magnitude of decline in metabolism is a strong predictor of the severity of the subsequent symptoms that develop.

Grey- and white-matter degeneration provide a second measure of non-specific neurodegeneration observed in individuals with AD using magnetic resonance imaging (MRI) ^8, 24, 27, 65, 66^. In contrast to hypometabolism, structural atrophy begins approximately five years prior to AD symptom onset ^8, 27, 67, 68^. While spatial patterns of grey matter atrophy for sporadic and autosomal dominant AD are similar, there are unique topological characteristics associated with each AD form ^33, 69, 70^. For example, ADAD individuals show atrophy in the putamen and thalamus, two regions not typically thought to undergo neurodegeneration in sporadic AD ^65^. These differences in grey-matter volumetric patterns further highlight the strength of using ADAD models, given this cohort allows for systematic, controlled, longitudinal studies of volumetric decline. Reductions in grey matter volume and thickness likely reflect differences in regional patterns of Aβ accumulation, evidenced by research showing that these same regions are known sites of increased PiB uptake specific to ADAD individuals ^8^.

In addition to the loss of grey matter, the integrity of white matter structures is also compromised in ADAD ^71–75^. This loss of white matter integrity is thought to reflect damage that occurs to neurons as a result of vascular changes related to Aβ deposits ^73, 76^ or tau protein hyperphosphorylation ^77^. In individuals carrying ADAD mutations, these integrity reductions have been reported globally, in the corpus callosum, and fronto-occipital fasciculus, as well as in the cingulum and fornix, two tracts that provide the input and output to the hippocampus respectively and potentially could contribute to the memory-deficits associated with cognitive symptom onset. Furthermore, within ADAD individuals, white matter disruption has been observed up to 10 years prior to symptom onset, particularly in the forceps major ^73^. These corpus callosal alterations are thought to potentially underpin functional network disruptions to the default mode network also observed in ADAD individuals ^78^. Importantly, posterior white matter alterations have been noted immediately prior to conversion from asymptomatic to symptomatic status in individuals outside of DIAN with a genetic predisposition to ADAD ^79^. In line with these reports, postmortem studies of AD have also indicated substantial myelin loss within these structures, as well as increased axonal degeneration in individuals with AD ^80^.

The DIAN Observational Study is a multi-site, longitudinal, study investigating preclinical ADAD using a combination of neuroimaging and other biomarker sampling methods. The study primarily assesses the preclinical phase of ADAD by comparing mutation carrying and non-carrying siblings from families with a known ADAD mutation. Individuals recruited into DIAN complete multimodal MRI, PET, biofluid sampling including CSF and blood, neuropsychological testing, and clinical assessments allowing for complex assessments of the trajectory of changes in these AD-biomarkers prior to ADAD symptom onset. Elucidation of this preclinical phase allows for the planning and implementation of clinical trials aiming to test potential drugs that could be used for AD treatments or cures. Moreover, the DIAN Observational Study provides a much-needed evidence-base from which trial design, including sample size estimates can be derived.

These principles are exemplified by the establishment of the DIAN Trials Unit (DIAN-TU). The first two drug arms of the DIAN-TU have recently concluded, and results will be leveraged to continue to develop and test novel drugs and interventions for the treatment of AD. A major goal of the DIAN Observational Study is to establish collaborations across the globe in the pursuit of understanding AD. To this end, the current manuscript aims to describe the neuroimaging data collected via this collaboration, and outline the acquisition and pre-processing parameters, to facilitate easy access to this data for all AD researchers.

## Methods

The imaging protocols for the DIAN Observational Study contain complementary acquisitions chosen to represent the most sensitive measures for detecting and understanding preclinical AD-related pathology. During the planning phase, considerations were made to accommodate concerns around time constraints, generalizability, feasibility of automated analyses, and well-tested acquisition methods. A major strategic decision was made to not require participants to know their mutation status, and therefore, at-risk individuals were included in the DIAN study. Throughout the course of the DIAN study, it has remained of paramount importance to not inadvertently reveal mutation status to participants who have not chosen to know their status. Below, the finalized measures are described in detail. Given that a major aim of the DIAN Observational Study was to create an open scientific resource, the resulting pre-processed data is available by request at https://dian.wustl.edu/our-research/observational-study/dian-observational-study-investigator-resources/. The remainder of this paper will outline the imaging data available in the most recent data release, DIAN Data Freeze 15, encompassing data collected from February 2008 through December 2020.

### DIAN Sites

The DIAN Observational Study was launched in 2008 with 10 sites. Since its inception, the study has grown to include a total of 22 centers that span the Americas, Australia, Asia and Europe (Figure 1, Figure 2). PET and MRI scanners at all sites are required to meet minimum hardware specifications to maximize our ability to harmonize data, and ensure equivalent image quality, across sites. All MRI scans were acquired on a 3 Tesla machine, while PET scans were acquired on one of the following PET scanner models: Siemens HR+, Siemens Biograph TruePoint PET/CT, Siemens Biograph mCT PET/CT, Siemens Biograph mMR, Siemens Biograph Vision PET/CT, Siemens High Resolution Research Tomograph (HRRT), Siemens Biograph 1023/1024, GE Discovery PET/CT, or Phillips PET scanner. Importantly, prior to any participants being recruited at each site, volunteer MRI scans were submitted to the Mayo Clinic (Rochester, MN) and volunteer PET scans were submitted to the University of Michigan teams for review to ensure each site’s hardware was able to produce images of sufficient quality in line with common Alzheimer Disease Neuroimaging Initiative (ADNI) protocols.

**Figure 1.**
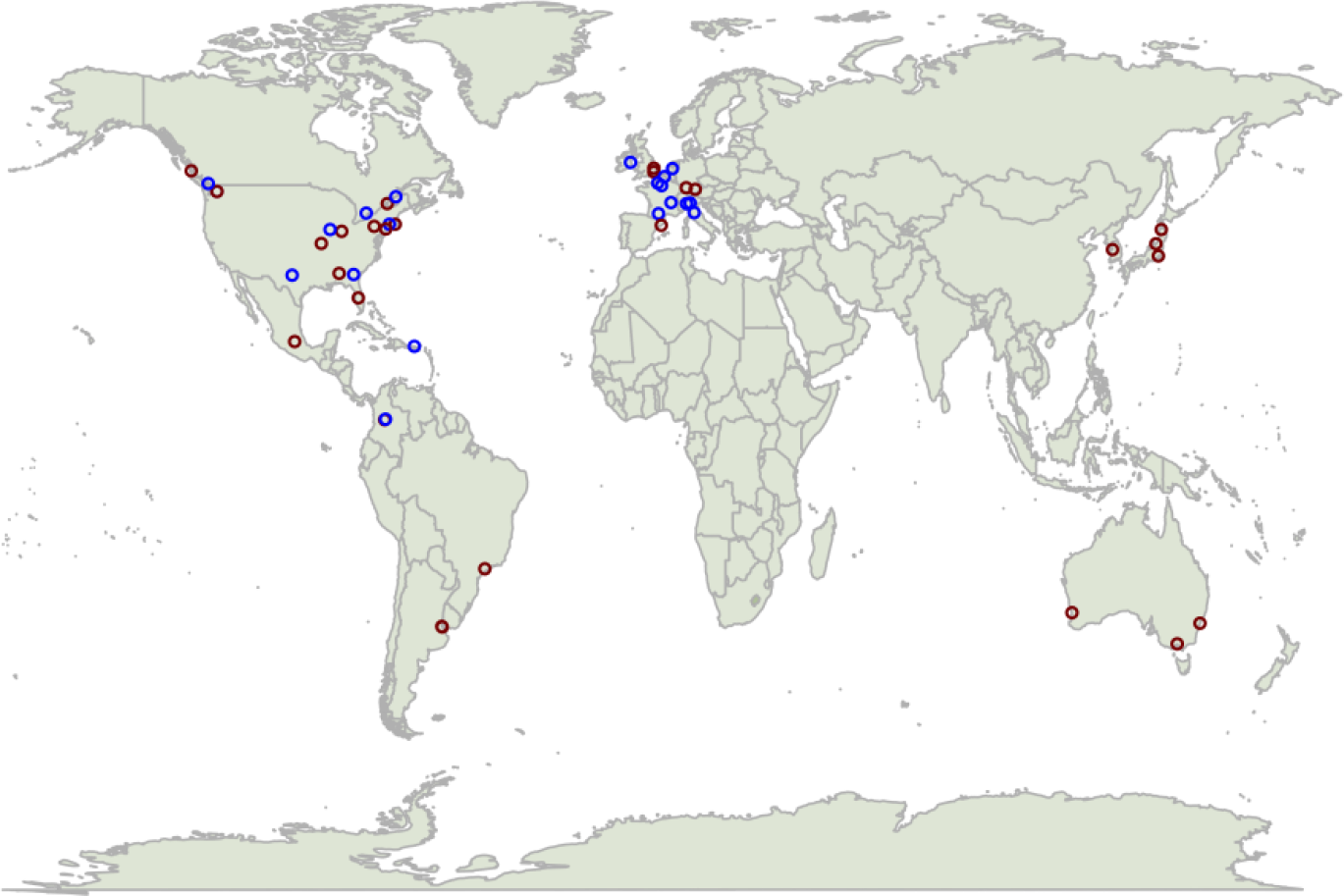
Participating DIAN sites as of 2021. Colors represent the arms of the DIAN study that each site is collecting data for. Blue dots represent sites collecting for the DIAN-Observational Study, red dots represent sites collecting for the DIAN-TU Study, and purple dots represent sites collecting for both DIAN-OBS and DIAN-TU. Together these depict a global network of AD researchers.

**Figure 2.**
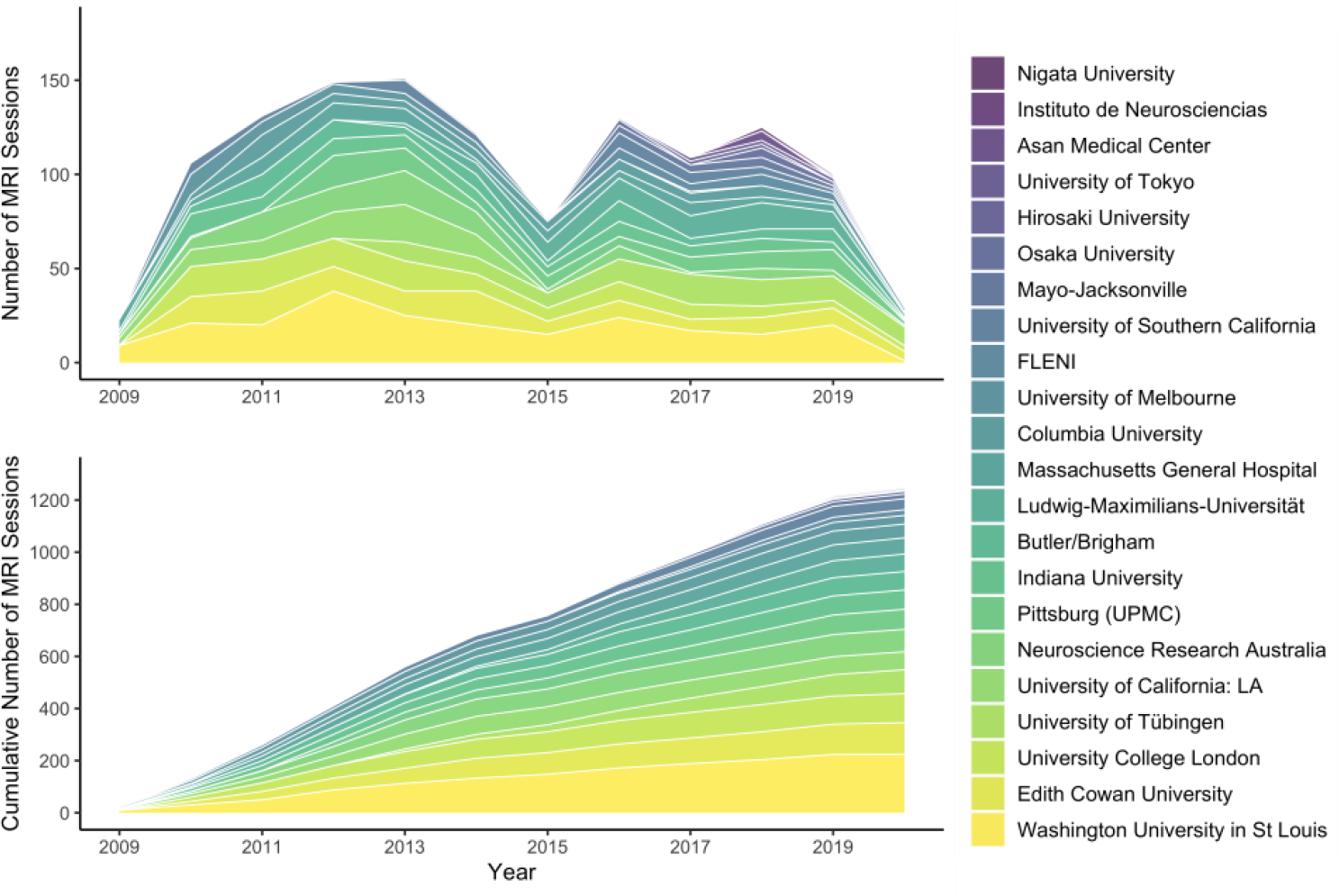
Stacked area plots depicting relative contributions to imaging data for each of the DIAN sites, and the overall evolution of this global study. At time of inception, DIAN consisted of 10 sites and grew to 22 active sites.

### Participants

Data from 533 participants, across 228 families with 114 different ADAD mutations spanning the *PSEN1*, *PSEN2*, and *APP* genes, are included in the most recent DIAN data freeze (DF15). These participants were recruited from DIAN performance sites, local initiatives at these sites, and through broader efforts (e.g., http://dian-i.org/, http://www.alzforum.org/new/detail.asp?id=1967, http://www.alz.org/trialmatch and http://www.dianexpandedregistry.org/). Importantly, the recruitment of offspring of a known ADAD mutation carrier results in participants having a 50% chance of inheriting the mutation that exists within their family. An individual’s mutation status is determined by genotyping, although participant mutation status is not revealed as part of the DIAN Observational Study. Independent genetic counselling and testing is made available to all participants. Mutation non-carriers are utilized as well-matched study controls for mutation carriers. As part of the DIAN Observational Study, participants were assessed on a battery of cognitive and clinical assessments every three years, unless the participant showed cognitive symptoms or was within three years of their expected symptom onset, in which case these tests were performed annually (Figure 3). All participants recruited gave informed consent to be included in the ongoing DIAN study.

**Figure 3.**
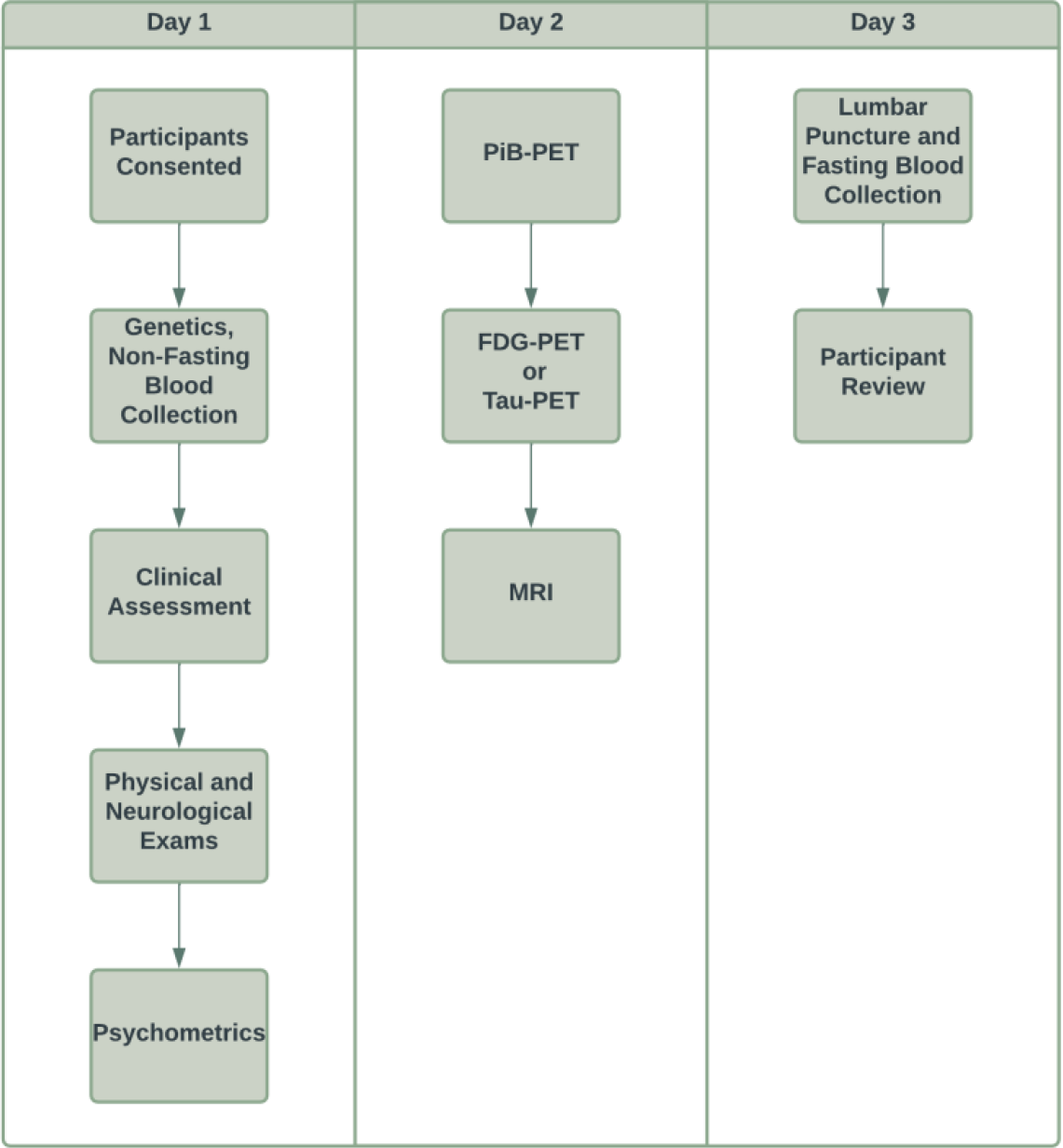
Chart depicting a typical DIAN research visit. In the original grant cycle, these visits were completed every three years, unless a participant is demonstrating cognitive symptoms, in which case they would begin to complete these assessments annually. After the DIAN grant cycle was renewed the visit frequency was updated to be completed every second year, unless a participant was symptomatic, in which case they would complete these visits yearly. Given most participants live large distances from the study sites, they typically travel to the location and complete all parts across three days. If participants do live in a study site city, they are able to complete the various components of the visit within three months. As the DIAN Observational Study begins to implement Tau-PET scans at applicable sites, these will be collected in place of the FDG-PET scans. Asymptomatic individuals stop in-person visits five years after they surpass their parental age of symptom onset, except in the case when they are known mutation carriers.

### Clinical Ratings

Participants were assessed using Clinical Dementia Rating^®^ (CDR^®^) scales to determine their dementia status ^81^. The overall CDR score was determined by evaluating CDR rating scales in memory, orientation, judgment, problem solving, function in community affairs, home and hobbies, and personal care. Using the resulting scores, individuals were classified as cognitively normal (CDR = 0), or having very mild (CDR = 0.5), mild (CDR = 1) or moderate-severe dementia (CDR > 1). In addition to CDR, participants were given a primary diagnosis as to the cause of any impairment.

Estimated years to onset (EYO) was assessed at each visit. EYO was calculated based on the participant’s current age, relative to their ‘mutation-specific’ expected age of dementia onset ^8^. The mutation-specific expected age of dementia onset was computed by averaging the reported age of dementia onset across individuals with the same mutation type. If the mutation-specific expected age at dementia onset was unknown, the EYO was calculated from the age at which parental cognitive decline began. The parental age of clinical symptom onset was determined by a semi-structured interview with the use of all available historical data provided by the participant or their caregiver. The EYO was calculated identically for both mutation-carriers and non-carriers. Importantly, all study staff performing clinical assessments were blinded to a participant’s mutation status.

### PET Acquisition

PET scans to measure Aβ deposition and brain glucose metabolism were performed across two days using a modified version of the ADNI PET acquisition protocol ^82^.

### PiB-PET

In order to measure Aβ deposition patterns, PiB-PET imaging was completed using a single bolus injection of approximately 15 mCi of [^11^C]PiB. PiB-PET scans were subsequently collected for either 70 minutes immediately following injection, or across a 30-minute time window that began after a 40-minute post-injection delay. For analyses, the common 30-minute period for each scan was used. Example PiB-PET images are displayed for each of the three participant group types in Figure 4. PiB-PET data has been used in many studies describing the deposition of amyloid within individuals who carry ADAD mutations ^33, 61^.

**Figure 4.**
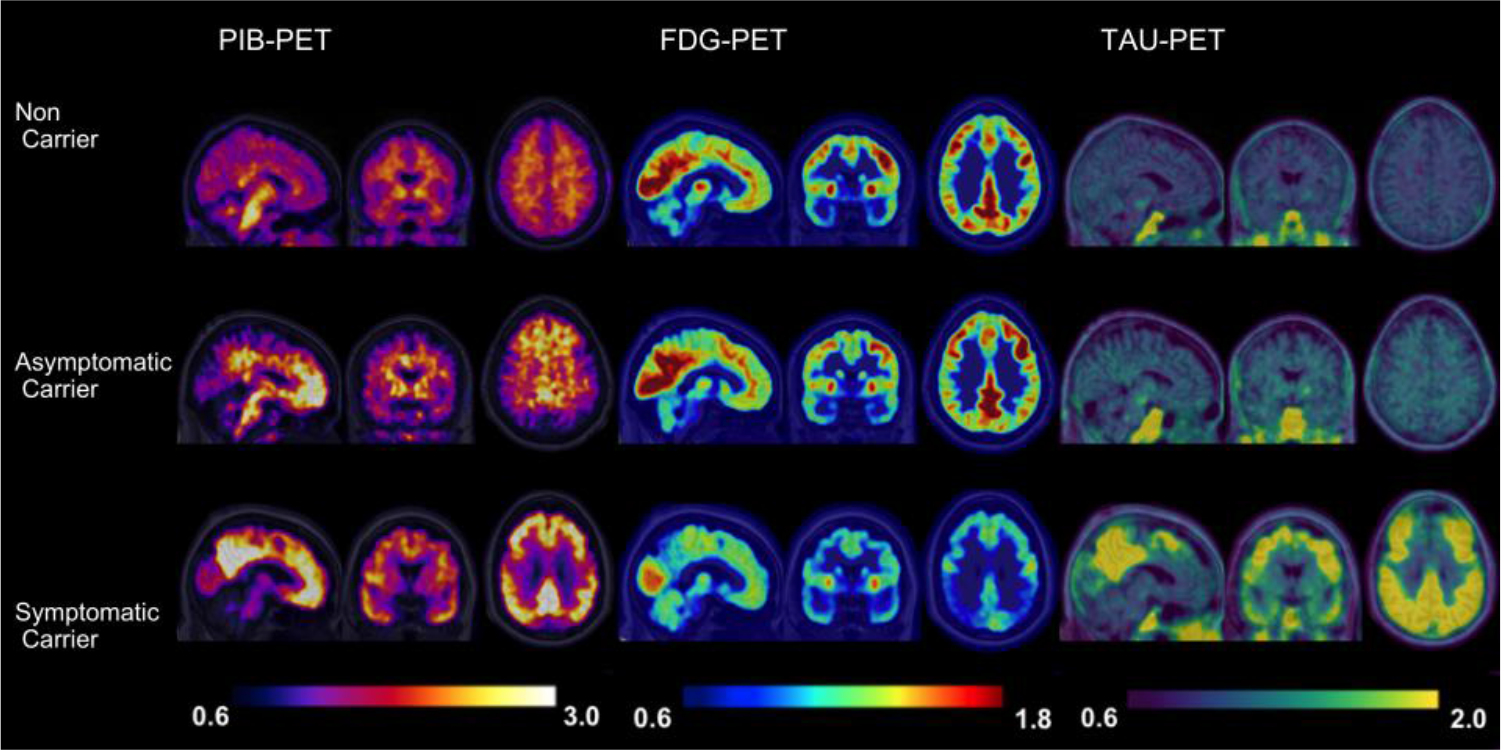
Example images representing each of the PET modalities collected across theDIAN study in a random participant from each group. **Left**, PIB-PET images depict PIB tracer uptake within an example individual who is a non-carrier (top row), asymptomatic mutation carrier (middle row), and symptomatic mutation carrier (bottom row). Higher levels of PIB tracer uptake is depicted in in both mutation carriers compared to the example non- carrier individual. A further increase of PIB tracer uptake is visible in the example symptomatic mutation carrier compared to the asymptomatic mutation carrier. These increases in PIB tracer indicate that these mutation carriers have increased Aβ deposition. **Middle** FDG-PET images depict FDG tracer uptake, where higher levels are associated with greater glucose metabolism. Here, we can visualize that the non-carrier (top row) and asymptomatic mutation carrier (middle row) example individuals have greater levels of glucose metabolism compared to the symptomatic mutation carrier (bottom row). **Right**, Tau- PET images depict AV-1451 tracer uptake. In both the example non-carrier (top row) and asymptomatic mutation carrier (middle row) we visualize relatively little AV-1451 uptake compared to the example symptomatic mutation carrier (bottom row). Higher AV-1451 uptake is associated with greater tau protein, which is expected in symptomatic stages of AD.

### FDG-PET

To measure glucose metabolism, approximately 5 mCi of [^18^F]FDG was given via a single bolus injection. Once the tracer was administered, a delay of 30 minutes was observed before the PET emission data were acquired for a period of 30 minutes. For analyses, the last 20-minute period of each scan was used. Example FDG-PET images are displayed for each of the three participant group types in Figure 4. To date, several studies have utilized the FDG-PET data to show differences in glucose hypometabolism for individuals with ADAD ^83–85^.

### Tau-PET

Given that the accumulation of tau has also been described as a characteristic pathology of AD ^33, 49^, tau-PET imaging is currently being added to the DIAN imaging protocol. In order to accommodate varying availability of tau-validated tracers across the globe, three tau-PET tracers are currently being added to the DIAN Observational Study: [^18^F]MK-6240, [^18^F]AV-1451 (flortaucipir), and [^18^F]PI-2620. At the point of writing, two of these tracers are actively being used by DIAN sites for data collection: [^18^F]MK-6240 and [^18^F]AV-1451 with the majority of the data being acquired with [^18^F]AV-1451. For [^18^F]MK-6240, a single bolus injection of 5 mCi of this tracer is given to participants, with images dynamically acquired for 110 minutes following injection, while for [^18^F]AV-1451, a single 10 mCi bolus injection of this tracer is administered to participants and images are acquired dynamically for the following 105 minutes. Standardized uptake value ratios (SUVrs) are calculated over the 80 -100 and 90 -110 minute post-injection windows for [^18^F]AV-1451 and [^18^F]MK-6240, respectively. Example tau-PET images for each of these two tracers in Figure 4.

### MRI Acquisition

Several MRI scan modalities were employed to visualize brain structure and function using MRI within the DIAN cohort. The order of scan collection was optimized to maximize participant compliance. For example, given participants are likely to move more as acquisition length increases, scans most critical for processing (i.e., T1-weighted) were run earlier within the protocol. All scans were collected using a 3 Tesla scanner, using parameters described below, although scanner model and manufacturer vary by site. All imaging modalities and their major goals are summarized in Table 1.

**Table 1.**
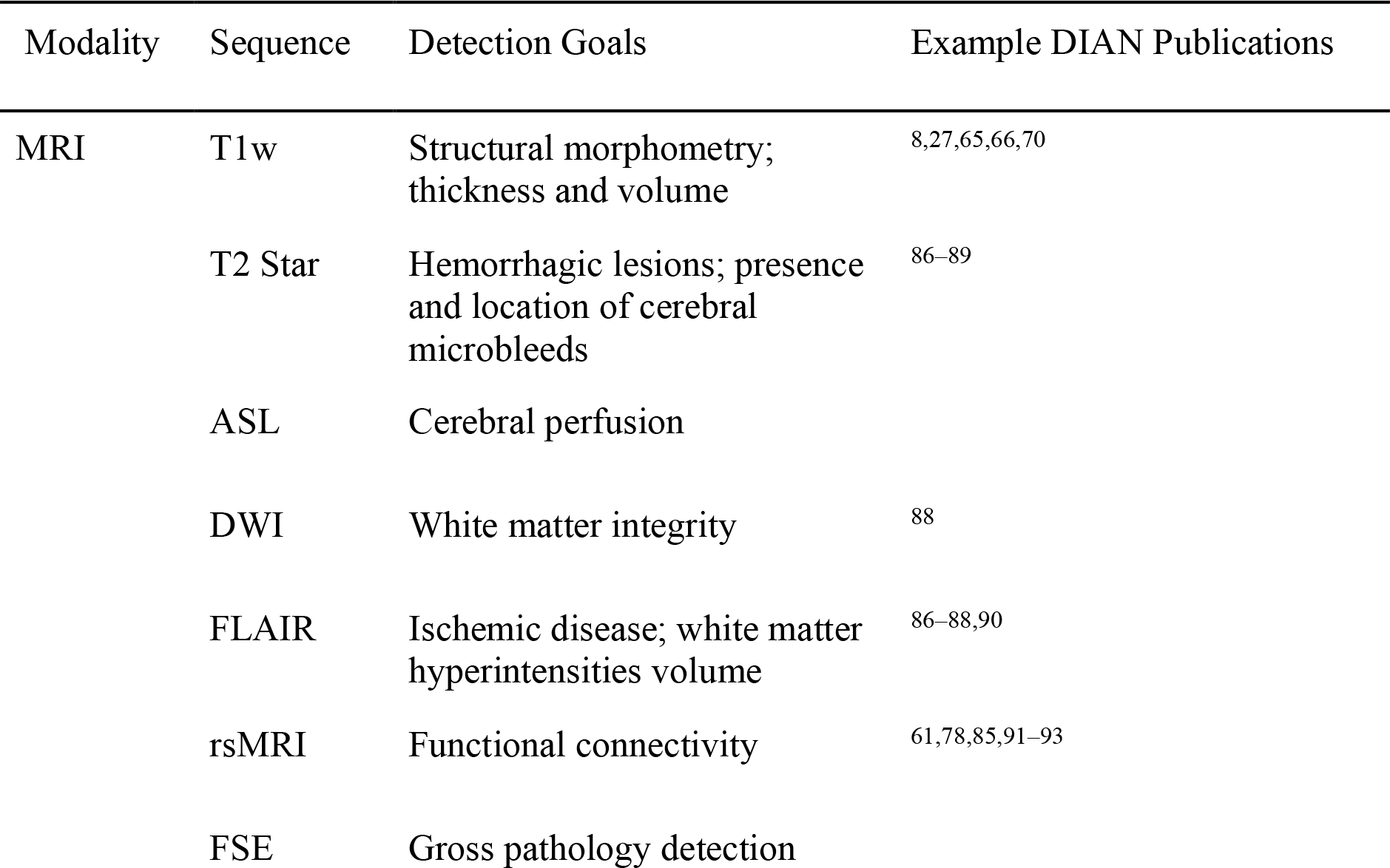
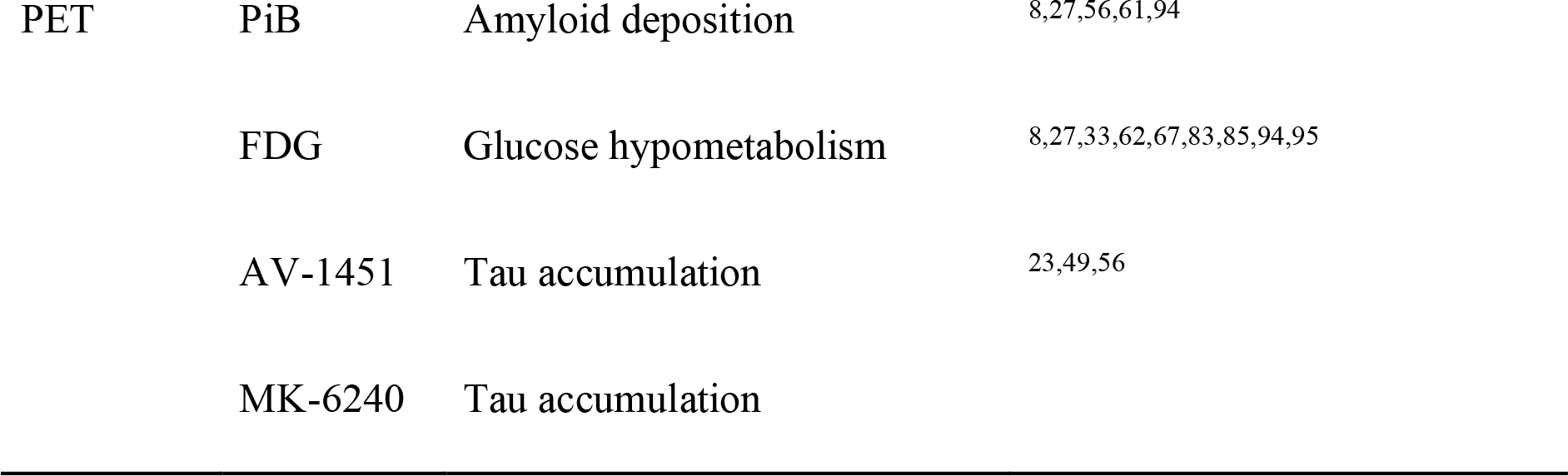
Overview of the Neuroimaging Modality Goals of the DIAN Observational Study, including key citations of published work involving these imaging modalities.

### T1-weighted Structural Scan

High quality T1-weighted scans providing information relating to brain volume and morphology are integrated into the pre-processing and registration of other collected scans (e.g., resting state MRI [rsMRI]), and are critical to the anatomical registration of PET data. The T1-weighted magnetized prepared rapidly acquired gradient echo (MPRAGE) sequence in DIAN was matched to the ADNI MRI protocol ^2, 82^, with the following parameters: TE = 2.95ms, TR = 2300ms, TI = 900ms, FOV = 270mm, flip angle = 9 degrees, number of slices = 225, voxel size = 1.1 x 1.1 x 1.2 mm^3^, GRAPPA acceleration factor = 2, acquisition time = 5 min 12 s. Example T1-weighted images are displayed for each of the three participant group types in Figure 5. To date, studies have utilized information derived from these high quality T1 images to determine structural patterns unique to ADAD ^24, 65, 66, 70^.

**Figure 5.**
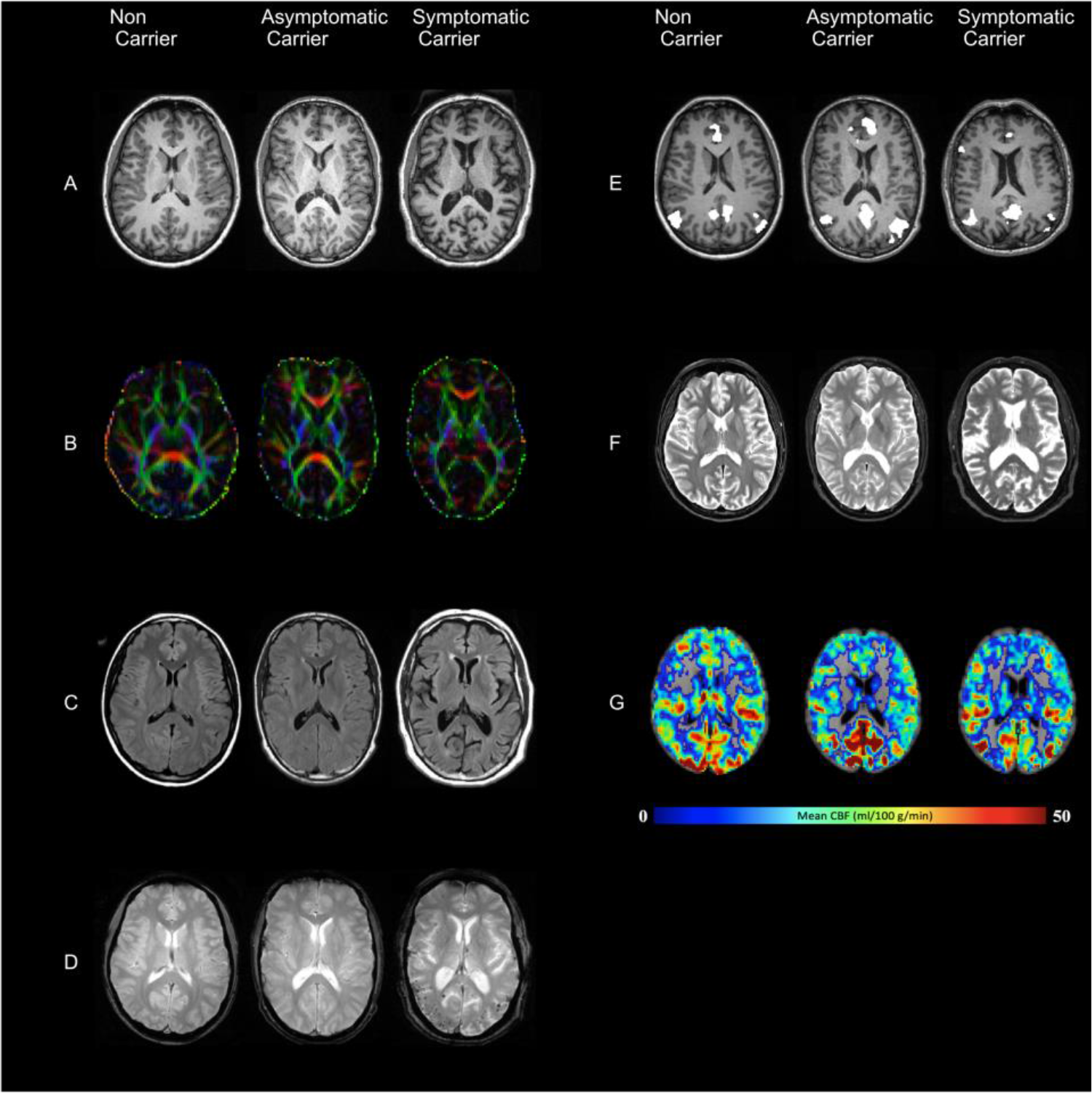
Example images representing each of the MRI modalities collected across the DIAN study in a random participant from each group. As briefly outlined in Table 1, each modality is collected to assess a specialized aspect of the brain. **A,** T1 images are collected to assess structural morphometry such as grey matter thickness and volume. Of note, ventricles in the example symptomatic carrier appear larger potentially signifying some atrophy in this individual compared to the example asymptomatic mutation carrier and non-carrier individuals. **B**, DTI images are collected in order to assess white matter integrity. Here, red, green, and blue, colors depict the primary direction of fiber orientation within each voxel, allowing researchers to assess microstructural changes in white matter integrity related to ADAD. **C**, FLAIR images are collected to assess white matter hyperintensities and edema. Here, in the symptomatic mutation carrier, several regions of white matter hyperintensities are visible as bright white lesions, compared to the example asymptomatic mutation carrier and non-carrier individuals. **D**, T2 STAR or susceptibility weighted images are collected to assess hemorrhagic lesions such as the presence and location of cerebral microbleeds, which are common in ADAD. In this example symptomatic mutation carrying individual, there are several cerebral microbleeds present, depicted as small black dots on the image. These microbleeds are not present in the asymptomatic mutation carrier, and non-carrier individuals. **E**, rsMRI images allow researchers to derive network representations to assess functional disruptions that may be present in association with ADAD disease stage. Here, the default mode network is revealed for each example individual by identifying regions of highly coherent activity fluctuations. **F**, FSE images are collected during this study in order to quickly assess large deviations from expected structural morphometry. For example, FSE images can be used to identify tumors or gross morphometrical issues such as extreme atrophy. **G**, ASL images are collected as a means of assessing cerebral perfusion. Researchers are able to derive maps of perfusion which allow them to quantify the rate of cerebral blood flow in any given individual. Lower perfusion and cerebral blood flow are associated with brain health and AD.

### Diffusion-weighted Imaging

Diffusion-weighted scans are specifically sensitive to the thermal motion of water over time, and therefore can be used to infer the presence of biological structures within the brain. Given the highly uniform nature of white matter within the brain, water movement tends to be especially constrained where axons are present ^96^. This property is not maintained within the cerebrospinal fluid (CSF) or grey matter structures, where water tends to diffuse in a much less constrained manner given there are relatively fewer biological boundaries in these structures ^97^. These properties of water movement make diffusion-weighted images uniquely positioned to measure the integrity of white matter microstructures within the brain. To this end, we initially acquired diffusion-weighted scans with the following parameters: TE = 81ms, TR = 7000ms, FOV = 256mm, number of slices = 60, voxel size = 2 x 2 x 2 mm^3^, maximum diffusion weighting = 2000s/mm^2^, acquisition time range = 2 min 43s – 3 min 04s. These traditional DTI-optimized scans were phased out in 2017 in favor of optimizing diffusion-weighted images for diffusion basis spectrum imaging (DBSI) in the DIAN-3 MRI protocol.

The DIAN-DBSI sequence comprises three diffusion sequence sessions with the Siemens built-in 6, 10, and 12 diffusion vectors, respectively. Multiple b-values were implemented in each session. The maximal b values for each session are 2000, 1500, and 1000 s/mm^2^, respectively. By combining all three sessions, a total of 28 unique directions were acquired, with 66 unique diffusion weightings. For each run, there was one volume with no diffusion weighting (b = 0 s/mm^2^) accounting for the remaining volumes. Together, the acquisition time was 9 min 14s. This unique sequence design allows for DBSI algorithms to subsequently implement neuroinflammation imaging, an important branch of diffusion research in preclinical AD (see Wang et al. ^98, 99^ for a full description of DBSI). Example diffusion-weighted images are displayed for each of the three participant group types in Figure 5.

### T2 FLAIR

T2 fluid-attenuated inversion recovery (FLAIR) images are useful tools for identifying white matter hyperintensities (WMH), a phenomenon known to be increased in individuals with sporadic- and autosomal dominant AD ^90, 100^. These scans complement diffusion-weighted images by providing information on the integrity of white mater macrostructure. Based on the ADNI MRI protocol, axial T2 FLAIR images were acquired with the following parameters: TE = 91ms, TR = 9000ms, TI = 2500ms, FOV = 220mm, flip angle = 150 degrees, slices = 35, voxel size = 0.9 x 0.9 x 5mm, acceleration factor = 2, acquisition time = 4m 05s. Example T2 FLAIR images are displayed for each of the three participant group types in Figure 5.

### T2 Star

T2 Star sequences can be used to characterize pathological changes occurring to venous vasculature within the brain. More specifically, this MRI sequence is sensitive to hemorrhage, calcification, and iron deposition ^101^, allowing researchers to detect the presence and location of cerebral microbleeds, a common pathology associated with ADAD ^77, 87–89^. Following the ADNI MRI protocol, we acquired axial T2 Star scans with the following parameters: TE = 20ms, TR = 650ms, FOV = 200mm, Flip angle = 20 degrees, slices = 44, voxel size = 0.8 x 0.8 x 4mm, acquisition time = 4m 11s. Example T2 Star images are displayed for each of the three participant group types in Figure 5.

### 2D or 3D Pulsed Arterial Spin Labelling

Arterial Spin Labeling (ASL) is an MRI technique that can measure blood perfusion without the use of an exogenous contrast agent and can be used to assess qualitative or quantitative cerebral blood flow (CBF). Previous studies using ASL have reported hypoperfusion in the posterior cingulate, precuneus, and parietal cortices in individuals with AD compared to healthy controls ^102–104^. However, few studies using ASL have focused on ADAD, and further investigations in this population are _needed_ 105,106. The multisite and international nature of DIAN necessitates use of readily available, standardized, and vendor provided ASL protocols, so protocols were harmonized to ADNI (adni.loni.usc.edu). Sequence parameters are provided in Table 2.

**Table 2.**
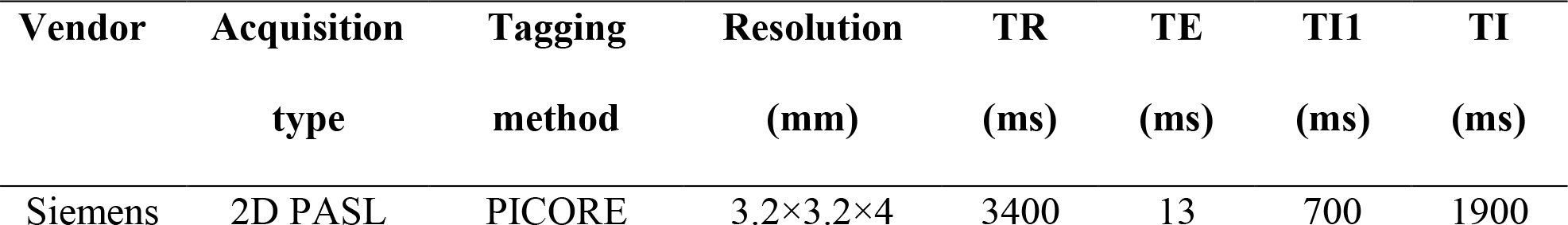
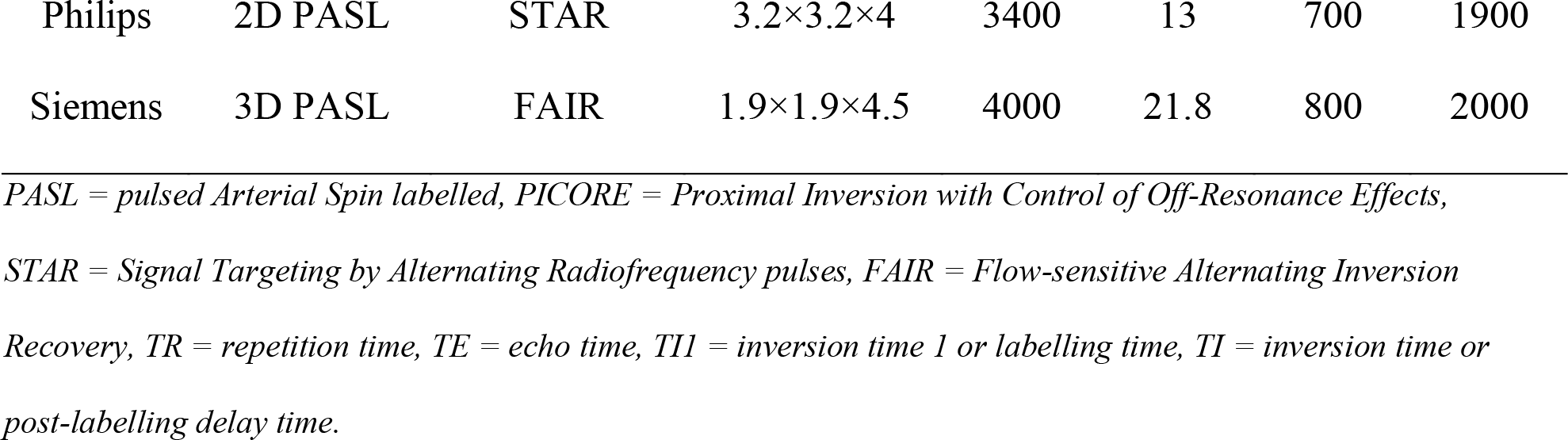
Parameters of the ASL sequence employed in the DIAN Observational study

| Vendor | Acquisition | Tagging | Resolution | TR | TE | TI1 | TI |
| --- | --- | --- | --- | --- | --- | --- | --- |
|  | type | method | (mm) | (ms) | (ms) | (ms) | (ms) |
| Siemens | 2D PASL | PICORE | 3.2×3.2×4 | 3400 | 13 | 700 | 1900 |
| Philips | 2D PASL | STAR | 3.2×3.2×4 | 3400 | 13 | 700 | 1900 |
| Siemens | 3D PASL | FAIR | 1.9×1.9×4.5 | 4000 | 21.8 | 800 | 2000 |

*PASL = pulsed Arterial Spin labelled, PICORE = Proximal Inversion with Control of Off-Resonance Effects, STAR = Signal Targeting by Alternating Radiofrequency pulses, FAIR = Flow-sensitive Alternating Inversion Recovery, TR = repetition time, TE = echo time, TII = inversion time 1 or labelling time, TI = inversion time or post-labelling delay time.*

### Resting State MRI

RsMRI scans can be employed to investigate low-frequency fluctuations in the blood oxygen level-dependent (BOLD) signal. These spontaneous changes in BOLD signal are thought to reflect intrinsic functional connectivity within the brain ^107^.

Previous work has shown that individuals with AD show abnormal patterns in rsMRI connectivity, particularly in the case of the default mode network ^108^. Given that this network is comprised of regions already implicated in AD pathology ^109^, rsMRI is a promising tool for helping researchers understand the connection between structural and functional disruption caused by ADAD progression ^61, 78, 91, 92^. Therefore, the following parameters were used to acquire the axial rsMRI scan: TE = 30ms, TR = 3000ms, FOV = 212mm, flip angle = 80 degrees, slices = 48, voxel size = 3.3 x 3.3 x 3.3mm, acquisition time = 5m 08s, eyes = open. This scan was repeated for a total acquisition time of 10m 16s. Example raw rsMRI images are displayed for each of the three participant group types in Figure 5.

### T2 FSE

In addition to the above scans, a T2 Fast Spin Echo (FSE) scan was also acquired as part of the MRI protocol. The main purpose of this scan is to assist in registration efforts for the rsMRI scans collected. Our T2 FSE scans were collected with the following parameters: TE = 563ms, TR = 3200ms, FOV = 270mm, slices = 225, voxel size = 1 x 1 x 1.2mm, GRAPPA acceleration factor = 2, acquisition time = 4m 08s. Example T2 FSE images are displayed for each of the three participant group types in Figure 5.

### Image Pre-processing and Quality Control

Once acquired, scans were screened for artefacts and compliance by the ADNI Imaging Core before being transferred to the pre-processing team. A standardized protocol was put in place for the transfer of raw data from the scanners to the site of pre-processing, Washington University in St Louis (Figure 6). Detailed MRI and PET protocol information beyond what is supplied in this manuscript is available on the study website, below.

**Figure 6.**
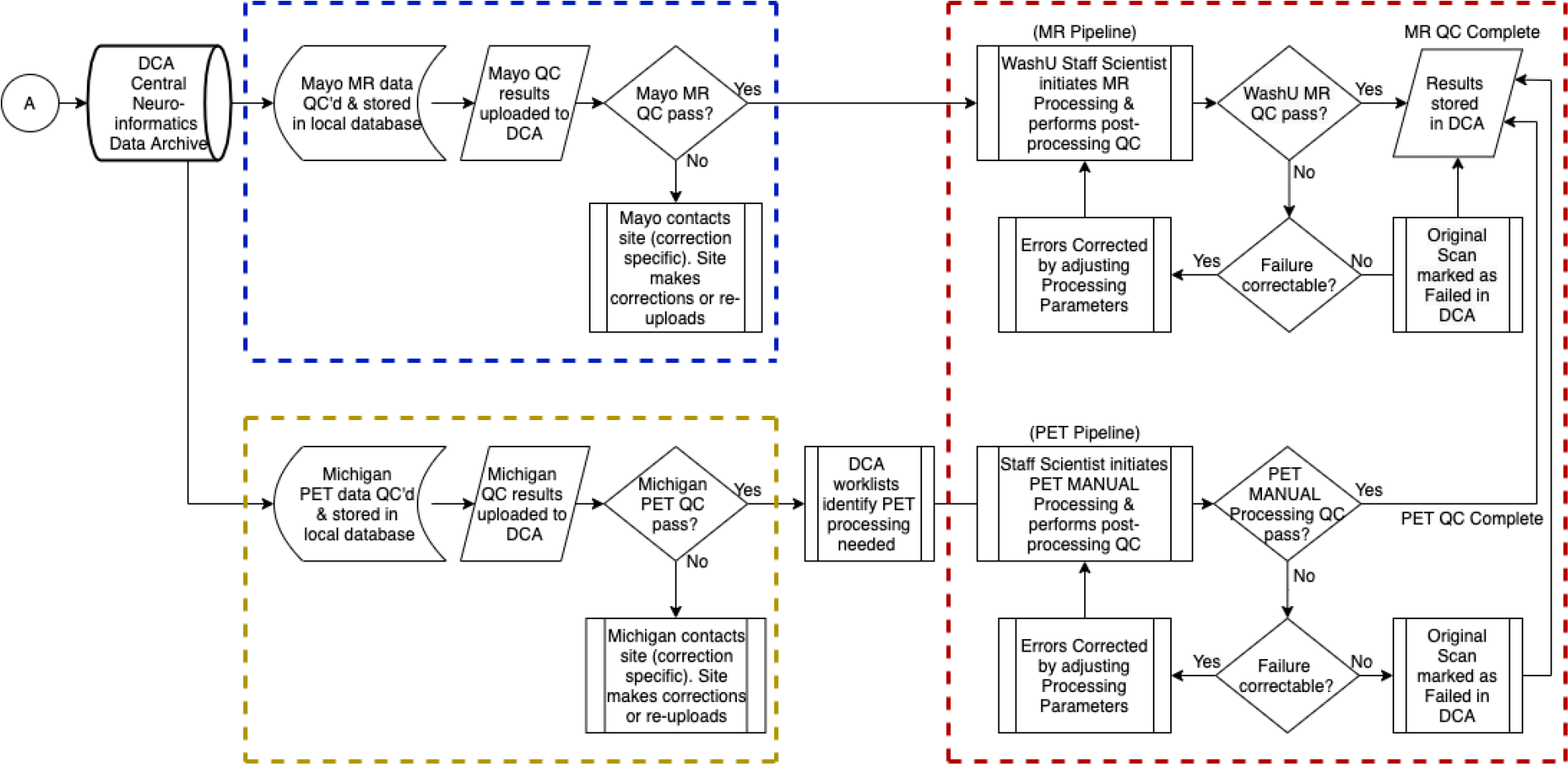
Flow chart depicting the quality control workflow for the DIAN Observational Study. Mayo Clinic, Rochester, MN, is responsible for the support, management, and primary quality control procedures for MRI, and for the participant safety reads (dashed blue lines). The University of Michigan, Ann Arbor, MI, is responsible for the support, management, and primary quality control of PiB- and FDG-PET participant sessions (dashed yellow lines). Once initial quality control has been passed, the MR and PET data are stored and processed in the DIAN Central Archive (DCA), an XNAT-based archive. Staff at Washington University School of Medicine are responsible for initial processing of MRI and PET images, and subsequently organizing data into each publicly accessible data release (dashed red lines).

### FreeSurfer

T1-weighted images were pre-processed using the FreeSurfer software suite (version 5.3-HCP-patch). In general, structural images were corrected for any artefacts ^110^, then a watershed and surface deformation procedure was used to remove the brain from the skull within the image ^111^. Images were then registered to Talairach space for subcortical white- and grey-matter structures to be segmented ^112, 113^. A tessellation step was then employed to estimate the grey and white matter structural boundary and apply any necessary topological correction ^114, 115^. All intersurface boundaries (white matter-grey matter, grey matter-cerebral spinal fluid) were placed in their optimal locations using surface normalization and intensity gradients ^116, 117^. Finally, images underwent surface inflation and registration to a spherical atlas ^118^. All images were manually inspected by a group of annually trained raters. Any images requiring manual intervention were corrected and the processing was rerun to ensure consistency across all scans. Up to three attempts were made to fix FreeSurfer errors that persisted after intervention. If errors continued to persist after the third attempt the FreeSurfer is considered to have failed quality control. Grey and white matter volumes, as well as cortical thickness for regions of interest in the Desikan atlas ^119^, were provided as the final output (for the full list of regions see: https://surfer.nmr.mgh.harvard.edu/fswiki/FsTutorial/AnatomicalROI/FreeSurferColorLUT).

### PET Quantification

PET data were analyzed using the PET Unified Pipeline (PUP; https://github.com/ysu001/PUP). PUP includes scanner resolution harmonization to a common full-width half-maximum ^120^, inter-frame motion correction using the summed image as the reference, PET to MR image registration, extraction of time activity curves for each FreeSurfer-defined region of interest (ROI) from the Desikan atlas, SUVr computations for each ROI, and a partial volume correction (PVC) procedure adjusting for regional spill-in and spill-out using a calculated regional spread function implemented in a geometric transfer matrix approach ^121, 122^. If dynamically acquired PET data are available (*n* = 324), PUP will additionally calculate non-displaceable binding potentials (BPND) using a Logan graphical analysis method ^123^. Quantitative modelling was performed on the post-injection time windows of 40 - 70 minutes and 40 - 60 minutes for PiB and FDG, respectively, with cerebellar grey chosen as the reference region.

## Results

### Overview

The current data set (DIAN Data Freeze 15) consists of information acquired from individuals who were recruited because their family had a known ADAD mutation. A total of 533 participants from 228 families have been enrolled. For this report, we have grouped participants into mutation non-carriers (*n* = 215), asymptomatic mutation-carriers *(n* = 214), and symptomatic mutation-carriers *(n* = 104). Four participants have been removed from further analyses because their genotype has not yet been determined. Within each group, there were equal proportions of males and females (*X^2^(2)* = 0.03, *p* = 0.98), left- and right-handedness (*X^2^(2)* = 0.69, *p* = 0.71), and no significant difference in *Apolipoprotein E* (APOE) allele frequencies (*X^2^(2)* = 0.76, *p* = 0.68). Importantly, for all subsequent analyses, a term has been added to statistical models to properly account for any shared variance that may exist amongst family members. However, there were significant differences for Age (*F(2,195)* = 59.11, *p <* 0.01), and number of years of education (*F(2,195)* = 13.7, *p <* 0.01). Symptomatic mutation-carriers had significantly higher average age (*M* = 45, *SD* = 10, *95%CI*: 41-49), and lower years of education (*M* = 13, *SD* = 3, *95%CI*: 12-14), compared to asymptomatic mutation-carriers (Age: *M* = 34, *SD* = 9, *95%CI*: 31-37; Education: *M* = 15, *SD* = 3, *95%CI*: 14-16), and non-carriers (Age: *M* = 37, *SD* = 11, *95%CI*: 36-38; Education: *M* = 15, *SD* = 3, *95%CI*: 14-15). To account for these differences, years of education and age were added to all subsequent models. Finally, symptomatic mutation-carriers were observed to have higher EYO at the initial visit (*M* = 3, *SD* = 5) compared to asymptomatic mutation-carriers (*M* = - 15, *SD* = 9) and non-carriers (*M* = -11, *SD* = 12). This result is unsurprising, given that the nature of their symptomatic status meant they were closer to symptom onset than other mutation-carriers. Given that mutation-carriers were split into asymptomatic and symptomatic sub-cohorts based on CDR, it does not make sense to analyze the proportion of CDR > 0 for these groups separately.

While these participants did complete a large number of cognitive tasks, the current paper focuses on the Mini Mental State Exam (MMSE) ^124^ and a general cognition composite score. This general cognition metric is derived from the average z-scored accuracy for the MMSE, Digit Symbol task of the Wechsler Adult Intelligence Scale-Revised battery ^125^, delayed logical memory subtask of the revised Wechsler Memory Scale ^126^, and delayed recall of the DIAN world list test. Together these tests index cognitive abilities such as verbal fluency, executive function, and memory, and were chosen to closely align with the planned cognitive composite of the DIAN-Trials Unit study ^42^. Symptomatic mutation-carriers had significantly poorer performance on the MMSE (*M* = 23.68, *SE* = 0.95, *95%CI*: 21.2-26.1) compared to asymptomatic mutation-carriers (*M* = 29.36, *SE* = 0.32, *95%CI*: 27.1-30) and non-carriers (*M* = 29.64, *SE* = 0.45, *95%CI*: 28.7-30; *F(2,193)* = 35.95, *p <* 0.01). Similarly, ymptomatic mutation-carriers had poorer average general cognition scores (*M* = -0.49, *SE* = 0.06, *95%CI*: -0.66- -0.31), indicating worse performance on a variety of cognitive tasks measuring memory and general cognition, compared to asymptomatic mutation-carriers (*M* = 0.13, *SE* = 0.03, *95%CI*: -0.03-0.28) and non-carriers (*M* = 0.14, *SE* = 0.03, *95%CI*: 0.07-0.20; *F(2,191)* = 109.6, *p* < 0.001). All demographic analyses are summarized in Table 3 and visualized in Figure 7.

**Figure 7.**
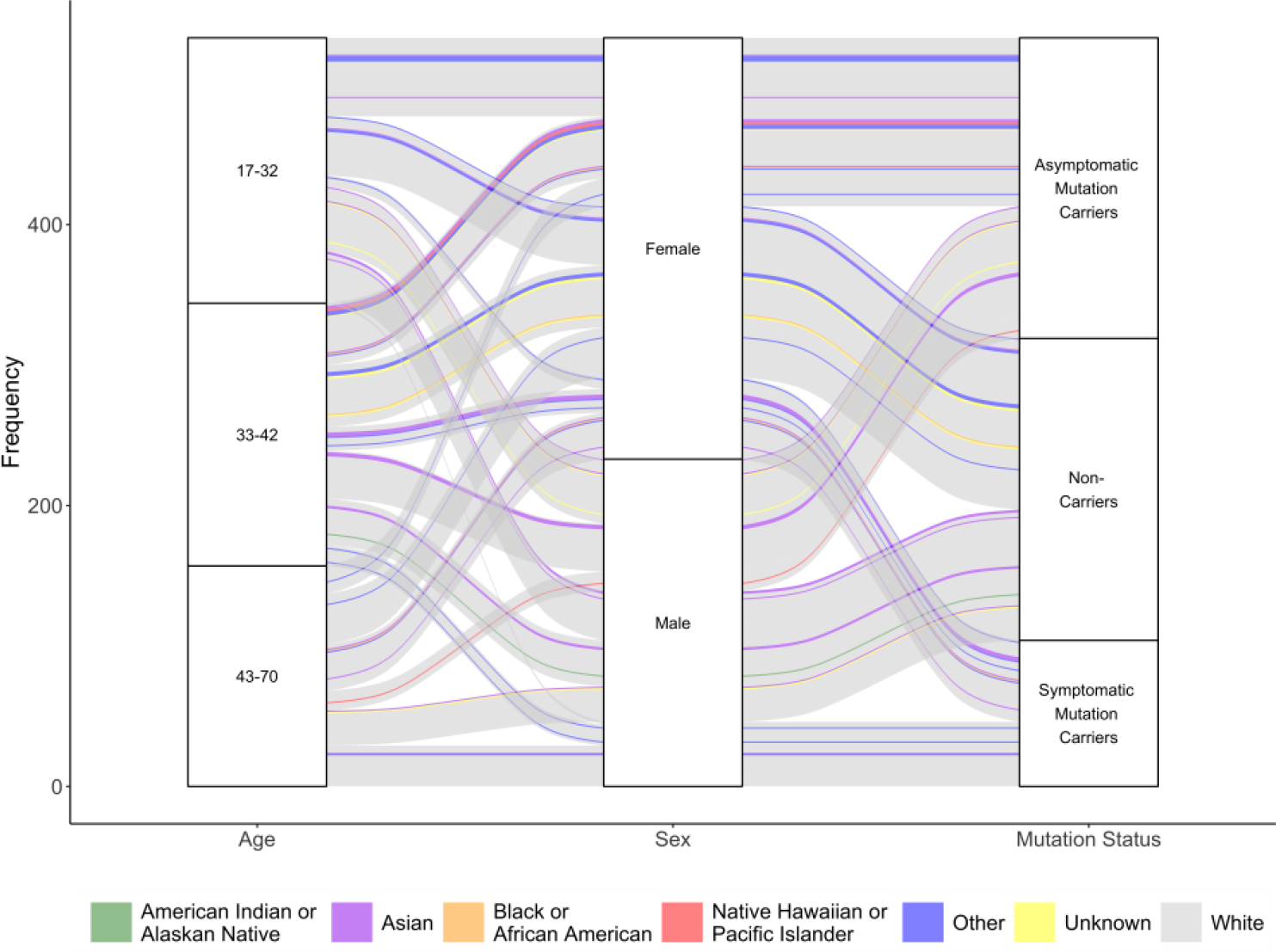
Schematic depiction of basic participant demographics. Age is displayed on the left panel, where each subdivision represents the proportion of participants that fall within that age bin. The middle panel represents the proportion of total participants within the DIAN study that identify as female and male. The final right-hand panel represents the proportion of individuals who are classified as non-carriers, asymptomatic mutation carriers and symptomatic mutation carriers. Racial composition of the DIAN cohort is represented in color, showing that while there are few individuals who identify as a race other than “white”, these individuals are present across all major demographic subdivisions of this cohort. Full breakdown of participants’ demographic information including self-reported racial identity. For race, individuals who selected “Other” self-identified as Arab (*n* = 2), Australian Aboriginal/Indigenous (*n* = 6), Croatian (*n* = 1), Hispanic (*n* = 3), Latin American (*n* = 3), Mexican American (*n* = 1), Middle Eastern (*n* = 2), Native Puerto Rican (*n* = 1), North African (*n* = 1), or none (*n* = 3), when provided as open-ended option.

**Table 3.**
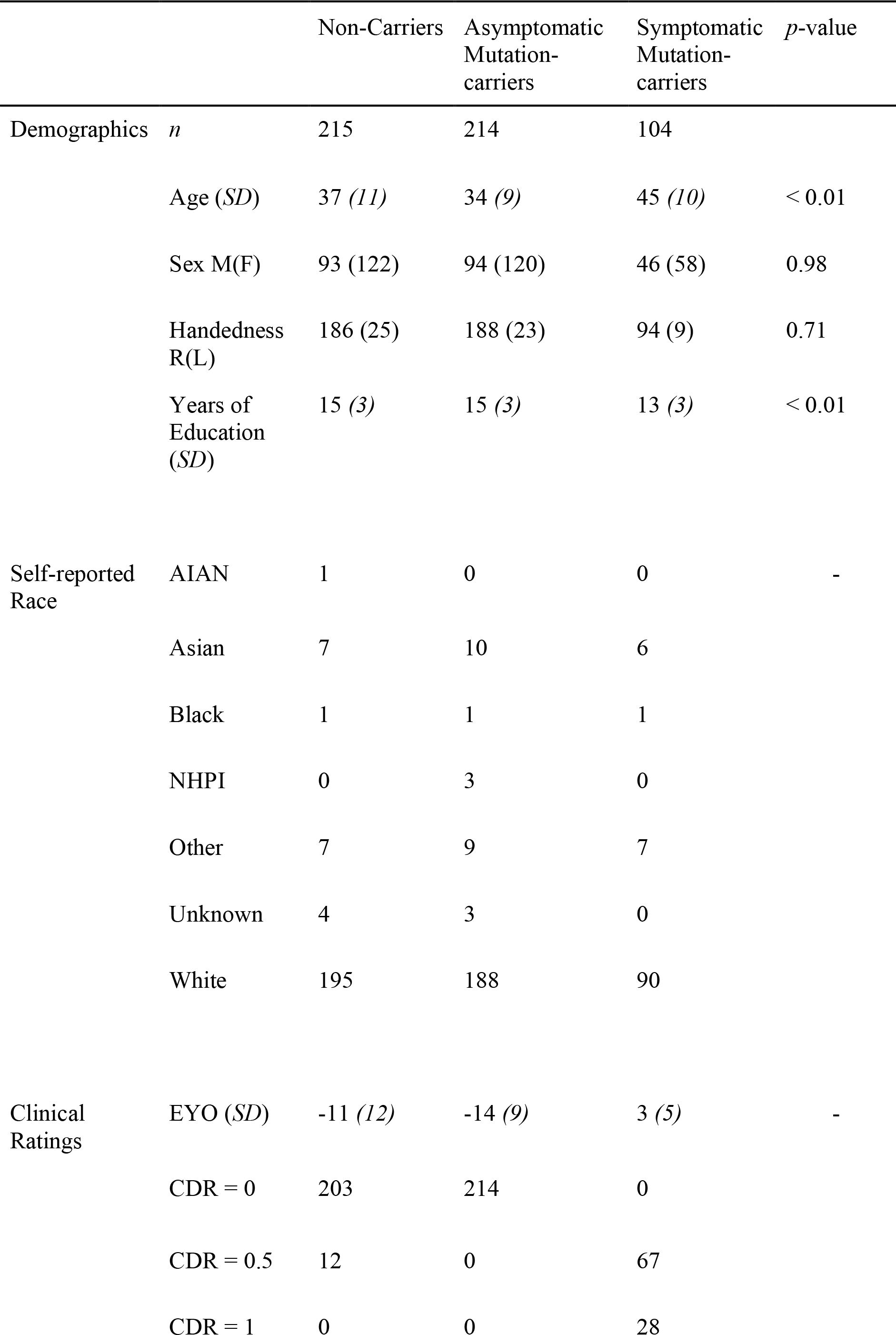
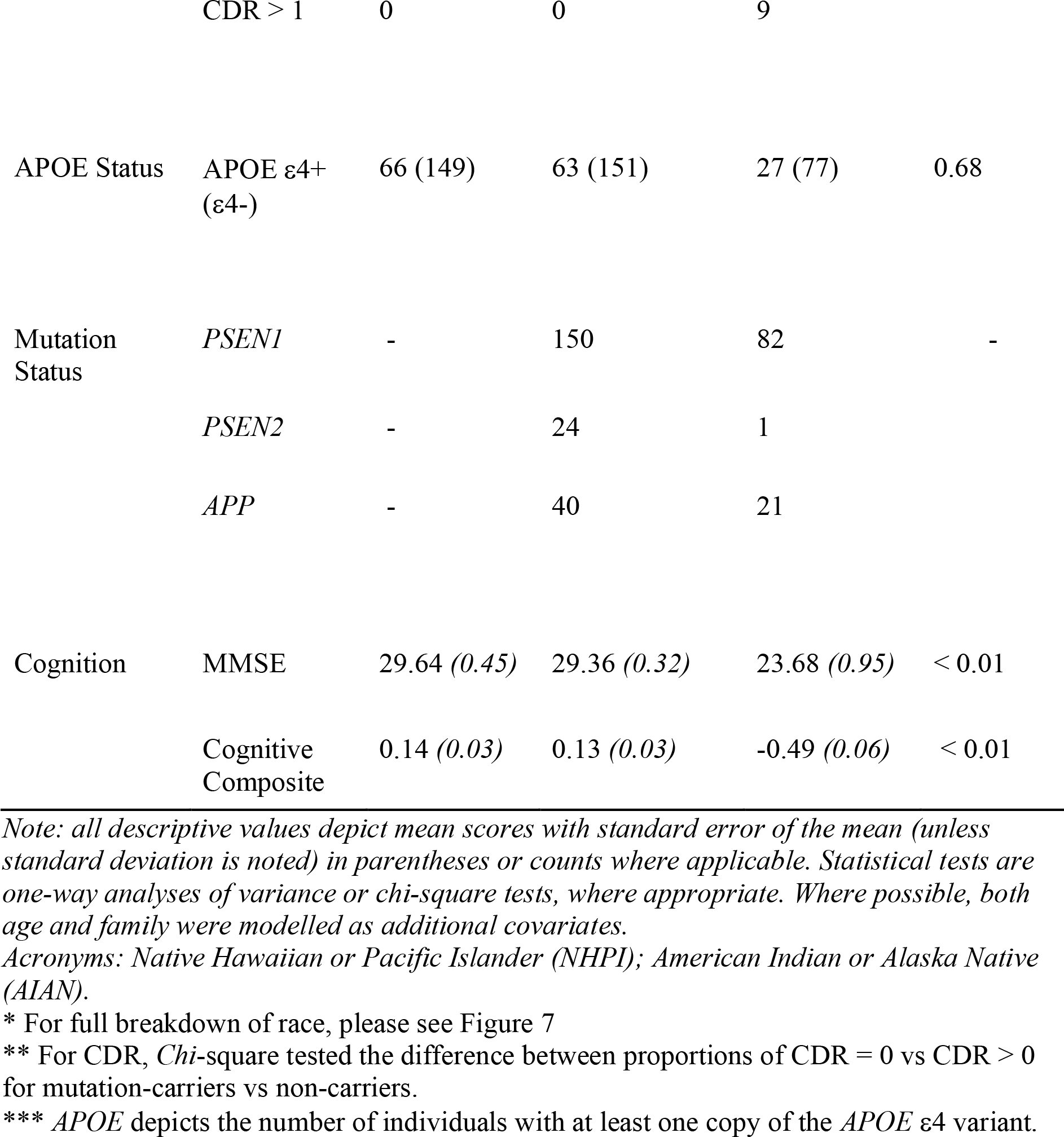
Demographic information for DIAN Data Freeze 15 Participants. Participants are classified by both their ADAD-mutation carrying status, as well as their score on the clinical dementia rating scale resulting in three independent groups: mutation non-carriers, asymptomatic mutation-carriers, and symptomatic mutation-carriers.

|  |  | Non-Carriers | Asymptomatic<br>Mutation-carriers | Symptomatic<br>Mutation-carriers | *p*-value |
| --- | --- | --- | --- | --- | --- |
| Demographics | *n* | 215 | 214 | 104 |  |
|  | Age (*SD*) | 37 (11) | 34 (9) | 45 (10) | < 0.01 |
|  | Sex M(F) | 93 (122) | 94 (120) | 46 (58) | 0.98 |
|  | Handedness<br>R(L) | 186 (25) | 188 (23) | 94 (9) | 0.71 |
|  | Years of<br>Education<br>(*SD*) | 15 (3) | 15 (3) | 13 (3) | < 0.01 |
| Self-reported<br>Race | AIAN | 1 | 0 | 0 | - |
|  | Asian | 7 | 10 | 6 |  |
|  | Black | 1 | 1 | 1 |  |
|  | NHPI | 0 | 3 | 0 |  |
|  | Other | 7 | 9 | 7 |  |
|  | Unknown | 4 | 3 | 0 |  |
|  | White | 195 | 188 | 90 |  |
| Clinical<br>Ratings | EYO (*SD*) | -11 (12) | -14 (9) | 3 (5) | - |
|  | CDR = 0 | 203 | 214 | 0 |  |
|  | CDR = 0.5 | 12 | 0 | 67 |  |
|  | CDR = 1 | 0 | 0 | 28 |  |
|  | CDR > 1 | 0 | 0 | 9 |  |
| APOE Status | APOE ε4+<br>(ε4-) | 66 (149) | 63 (151) | 27 (77) | 0.68 |
| Mutation<br>Status | *PSEN1* | - | 150 | 82 | - |
|  | *PSEN2* | - | 24 | 1 |  |
|  | *APP* | - | 40 | 21 |  |
| Cognition | MMSE | 29.64 (0.45) | 29.36 (0.32) | 23.68 (0.95) | < 0.01 |
|  | Cognitive<br>Composite | 0.14 (0.03) | 0.13 (0.03) | -0.49 (0.06) | < 0.01 |

*Note: all descriptive values depict mean scores with standard error of the mean (unless standard deviation is noted) in parentheses or counts where applicable. Statistical tests are one-way analyses of variance or chi-square tests, where appropriate. Where possible, both age and family were modelled as additional covariates.*
*Acronyms: Native Hawaiian or Pacific Islander (NHPI); American Indian or Alaska Native (AIAN).*
\* For full breakdown of race, please see Figure 7
\*\* For CDR, *Chi-square* tested the difference between proportions of CDR = 0 vs CDR > 0 for mutation-carriers vs non-carriers.
\*\*\* *APOE* depicts the number of individuals with at least one copy of the *APOE* ε4 variant.

### Scan Availability

Within the DIAN Observational Study, individuals completed three separate sessions of neuroimaging scans: MRI, PiB-PET, and FDG-PET. Given that some individuals did not complete all sessions, or their scans might have failed QC assessments, Table 4 summarizes the number of scans that are available within the DIAN Data Freeze 15, by group.

**Table 4.**
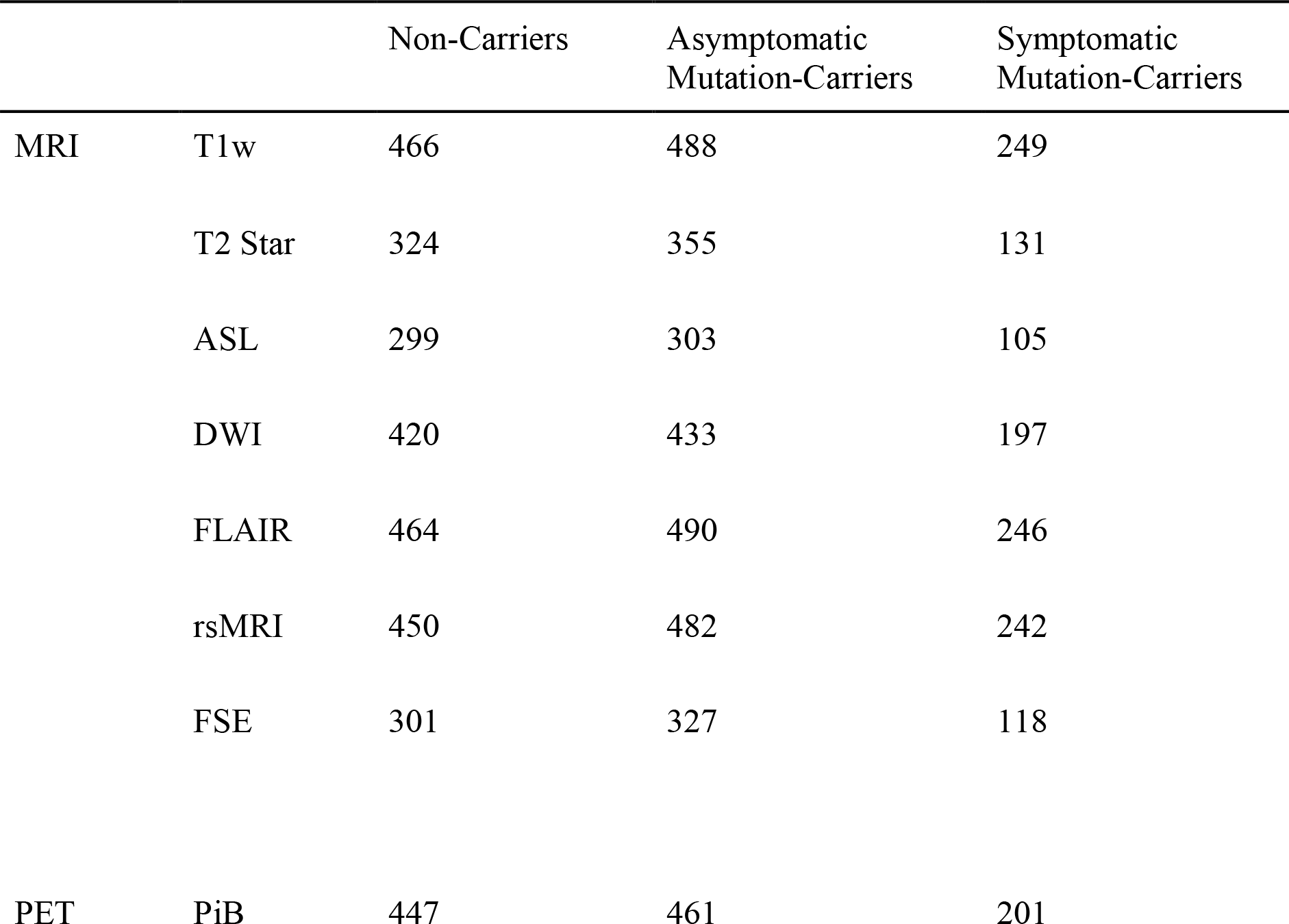
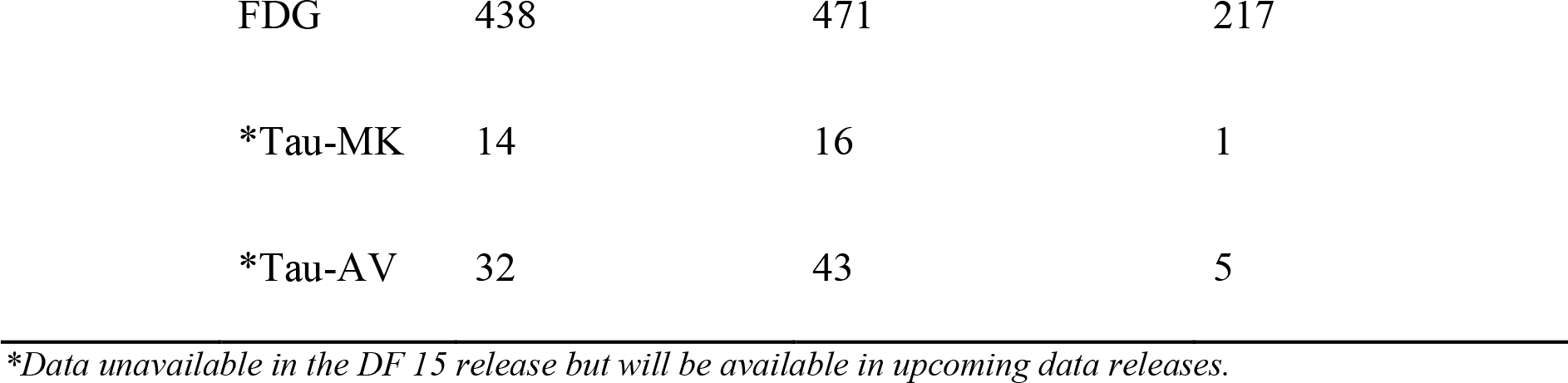
Total number of each scan type available in Data Freeze 15. Note, Tau-MK represents images acquired using the Tau-Pet tracer MK-6240, while Tau-AV represents images acquired using the AV-1451 tracer.

Furthermore, Table 5 outlines the numbers of individuals who have completed multiple visits as part of the longitudinal aims of the DIAN Observational Study. Example scans are depicted for each major structural mode measured within this cohort, below.

**Table 5.**
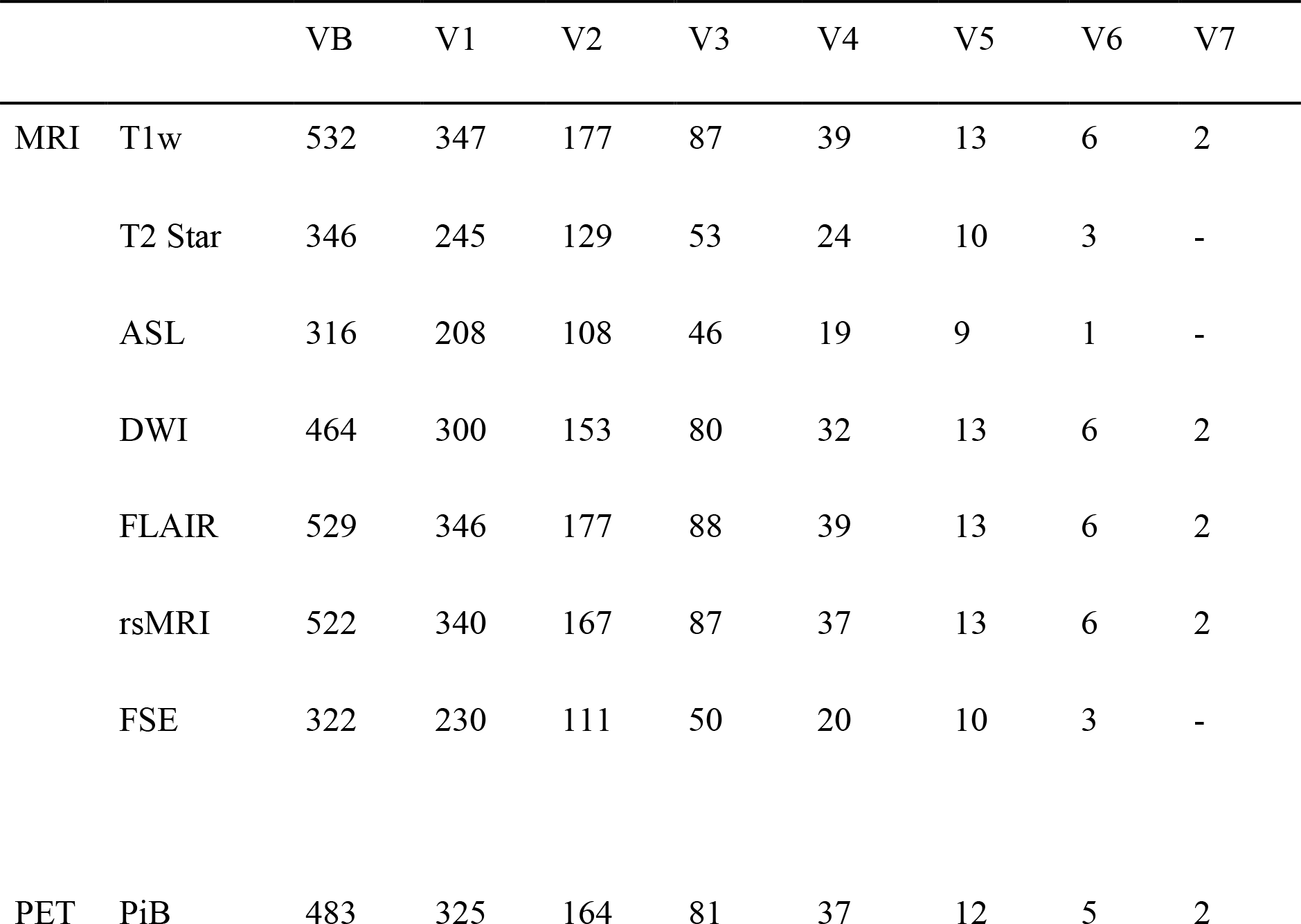
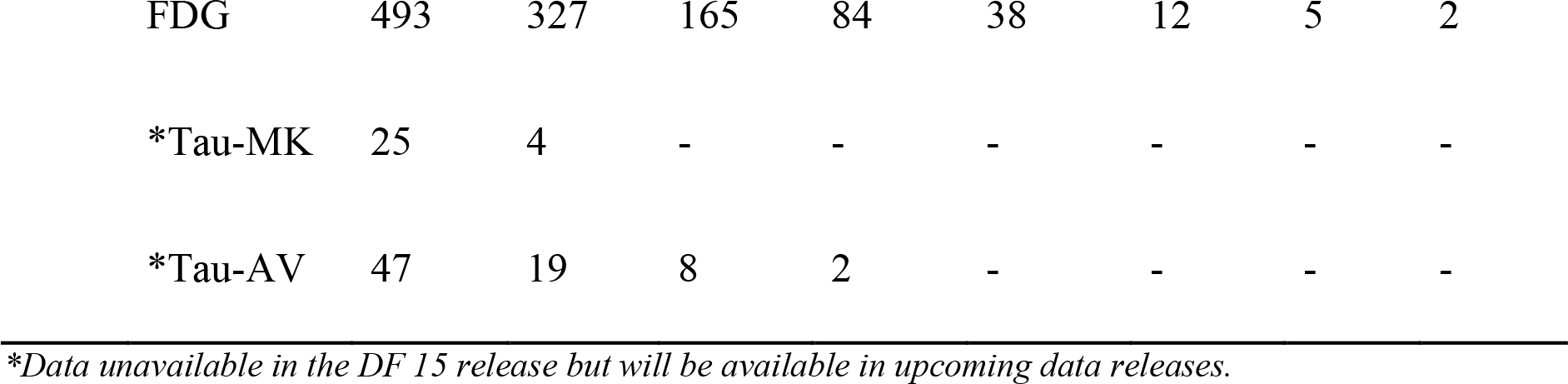
Longitudinal availability of images by scan type. Here VB depicts baseline imaging visit, and V1-V7 depict the subsequent imaging visits. Note, Tau-MK represents images acquired using the Tau-Pet tracer MK-6240, while Tau-AV represents images acquired using the AV-1451 tracer. For all statistical comparisons, data collected at VB is used, hence, this is the sample size for each independent modality.

### Image Analyses

Analyses of SUVr summary values for each of the PiB-PET and FDG-PET scans show differences in SUVr for symptomatic mutation-carriers compared to asymptomatic mutation-carriers and non-carriers. A summary AD-specific cortical PiB-PET measure was computed as the average SUVr of the FreeSurfer-Desikan atlas-derived lateral orbitofrontal, mesial orbitofrontal, rostral mesial frontal, superior frontal, superior temporal, mesial temporal, and precuneus regions ^127^. Using this measure, symptomatic mutation-carriers were observed to have significantly higher PiB-PET SUVr signifying increased Aβ deposition (*M* = 2.59, *SE* = 0.16, *95%CI*: 2.26-2.93) compared to both asymptomatic mutation-carriers (*M* = 1.57, *SE* = 0.07, *95%CI*: 1.28-1.85), and non-carriers (*M* = 1.07, *SE* = 0.02, *95%CI*: 0.95-1.18; *F(2,103)* = 237.70, *p* < 0.01). Similarly, a summary AD-specific FDG-PET measure was computed as the average SUVr of the FreeSurfer-Desikan atlas-derived isthmus cingulate and inferior parietal regions based on previous work outlining these are regions of interest in FDG-PET studies ^128, 129^. Symptomatic mutation-carriers were found to have significantly lower FDG-PET SUVr for this measure (*M* = 1.47, *SE* = 0.02, *95%CI*: 1.40-1.53) indicating lower levels of glucose metabolism compared to asymptomatic mutation-carriers (*M* = 1.68, *SE* = 0.01, *95%CI*: 1.62-1.73), and non-carriers (*M* = 1.71, *SE* = 0.01, *95%CI*: 1.68-1.73; *F(2,177)* = 120.28, *p* < 0.01). All PET analyses are summarized in Table 6 and Figure 8, representative PET scans are depicted in Figures 4.

**Figure 8.**
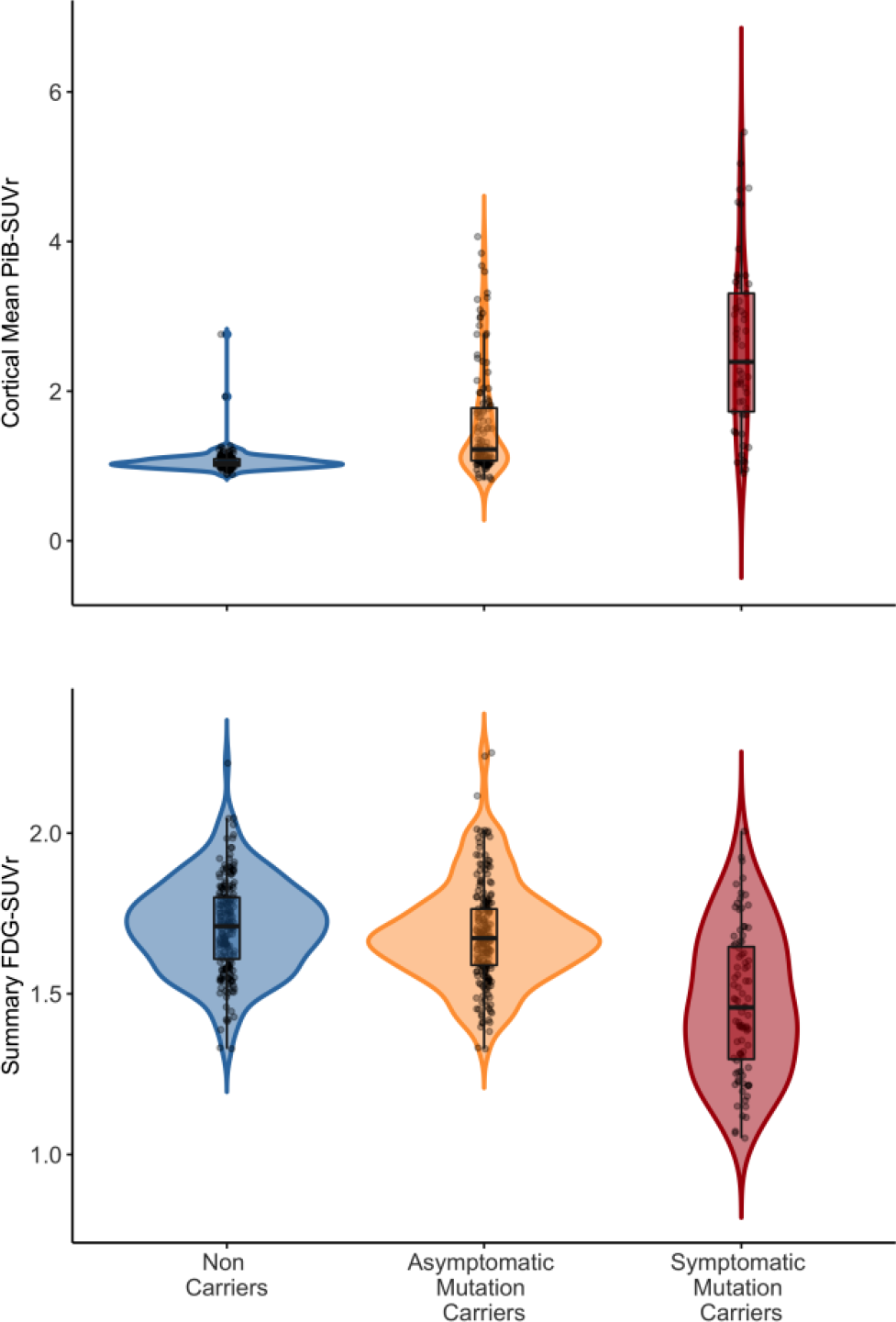
Summary depiction of the results of the PiB- and FDG-PET analyses. Violins represent the overall distribution of SUVr uptake for each tracer, for each group of participant groups, while box plots show the interquartile ranges for these scores. The horizontal thick black line depicts the median SUVr value for each group, while the whiskers depict the 95% confidence intervals. Depicted in these plots, symptomatic mutation carriers appear to have increased hypometabolism and amyloid tracer uptake, indicative of increased ADAD-related pathology. Using all available data, all FDG analyses consist of 197 non-carriers, 205 asymptomatic mutation carriers and 91 symptomatic mutation carriers, while the PIB analyses consist of 130 non-carriers, 120 asymptomatic mutation carriers, and 53 symptomatic mutation carriers.

**Table 6.**
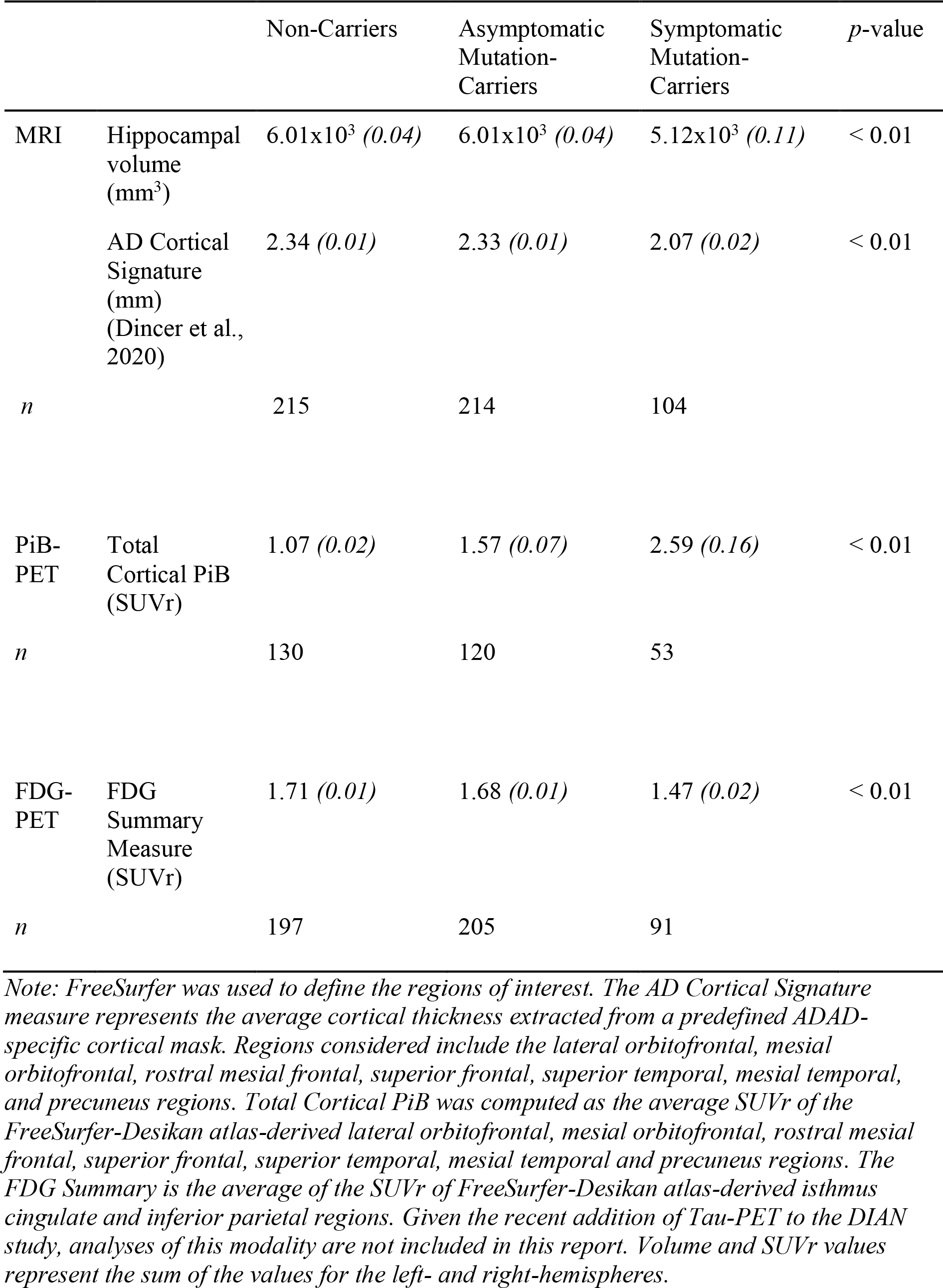
Summary results of the PET and MRI comparisons of the three participant groups that make up the DIAN study.

Additionally, analyses of T1-weighted MRI data pre-processed using FreeSurfer show that symptomatic mutation-carriers have significantly smaller total hippocampal volumes during their baseline visit (*M* = 5.12x10^3^, *SE* = 0.11x10^3^*, 95%CI*: 4.8-5.4) compared to asymptomatic mutation-carriers (*M* = 6.01x10^3^, *SE* = 0.04x10^3^*, 95%CI*: 5.8-6.2), and non-carriers (*M* = 6.01x10^3^, *SE* = 0.04x10^3^*, 95%CI*: 5.9-6.1; *F(2,193)* = 90.52, *p* < 0.01).

Furthermore, we were interested in using the cortical signature mask outlined in Dincer et al. (2020), in order to determine whether mutation-carriers differed in cortical thickness in regions known to be specifically linked to decline in AD. Extracting thickness measures using this mask, we found decreased total thickness within these AD-specific regions for symptomatic mutation-carriers (*M* = 2.07, *SE* = 0.02, *95%CI*: 2.0-2.1) compared to asymptomatic mutation-carriers (*M* = 2.33, *SE* = 0.01, *95%CI*: 2.2-2.6), and non-carriers (*M* = 2.34, *SE* = 0.01, *95%CI*: 2.3-2.4; *F(2,193)* = 267.7, *p* < 0.01). All MRI analyses are summarized in Table 6 and Figure 9, representative MRI scans are depicted in Figures 5.

**Figure 9.**
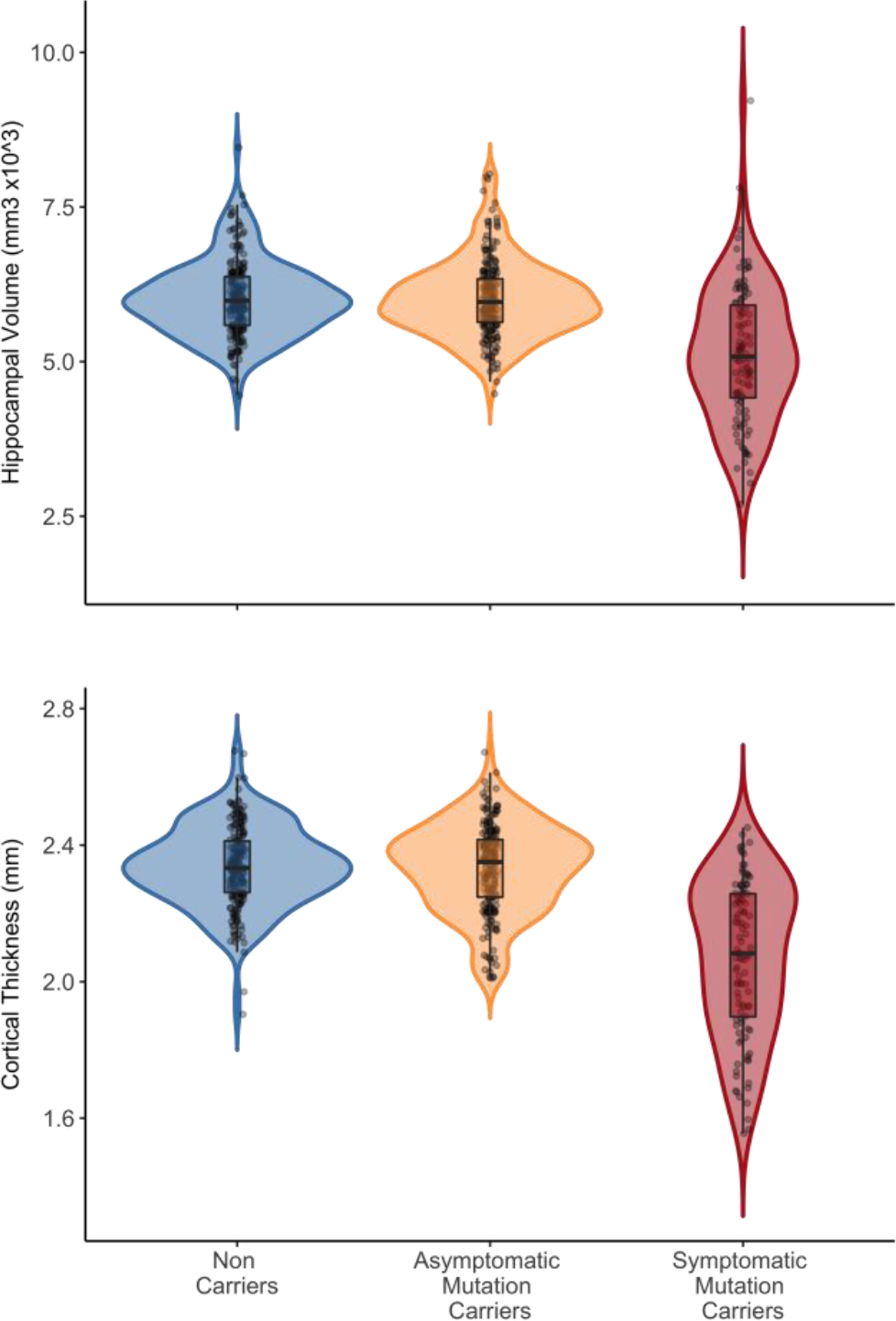
Summary depiction of the results of the analyses performed on the T1-weighted images. Violins represent the overall distribution of hippocampal volume and cortical thickness for each group, respectively. Similarly, box plots show the interquartile ranges for these scores, while the horizontal thick black line depicts the median value for each group, while the whiskers depict the 95% confidence intervals. In line with prior work, symptomatic mutation carriers have lower average cortical thickness and volume in ADAD-specific grey matter regions. Using all available scans, these analyses were conducted on 215 non-carriers, 214 asymptomatic mutation carriers, and 104 symptomatic mutation carriers.

## Discussion

The DIAN Observational Study contributes to our growing understanding of AD by facilitating the global study of individuals from families with pathogenic mutations in the *PSEN1*, *PSEN2*, and *APP* genes. Given that age at symptom onset is highly heritable within families and for individual mutations, the DIAN Observational Study uniquely allows researchers to study the preclinical phases of ADAD by staging participants based on their EYO. The mutations studied within the DIAN cohort are autosomal dominant, therefore, the offspring within these mutation-positive families have a 50% chance of inheriting the mutation from their affected parent. Individuals who do not inherit their family’s mutation are also included in the DIAN Observational Study as well-matched controls for environmental and other genetic factors that could be influencing disease expression and also as a true normal population. Together, these study design factors, along with the rich data collected across many biomarker fields, allow the DIAN Observational Study to investigate aspects of ADAD with higher confidence than has been possible for the study of sporadic AD. To that end, the DIAN Imaging Core Data Freeze 15 has collected and processed 1203 high quality T1-weighted scans, as well as a similar number of T2 STAR, T2 FLAIR, Diffusion, ASL, rsMRI, PiB-PET, and FDG-PET scans. These data are freely available upon request and represent the first resource of this magnitude covering a diverse range of ADAD biomarker data.

Results presented above describe example scans from the PET and MRI protocols employed as part of the DIAN Observational Study. This manuscript does not intend to comprehensively analyze these data, and instead hopes to highlight some basic characteristics of the data. Our PET and MRI results are consistent with previous work demonstrating that neuropathology, brain atrophy, and functional disturbance can be detected in the preclinical phase of AD ^7, 8^. At baseline visit we observed that symptomatic mutation-carriers have higher PiB uptake indicating greater Aβ deposition compared to asymptomatic mutation-carriers and non-carriers; a pattern that has previously been described in the broader literature ^130–133^. Using FDG-PET, our analyses show that symptomatic mutation-carriers have disturbed cortical glucose metabolism compared to asymptomatic mutation-carriers and non-carriers. These results are congruent with previous reports of global hypometabolism in preclinical AD ^134, 135^, as well as reports of hypometabolism in this group specific to the parieto-occipital regions ^27^. Finally, MRI analyses using structural images show that symptomatic mutation-carriers have reduced hippocampal volume and reduced cortical thickness in regions thought to be specifically impacted by AD, compared to asymptomatic mutation-carriers and non-carriers. This result is supported by other studies also highlighting hippocampal volumetric differences ^8, 65, 136^, and reduced thickness ^70, 137–139^ in individuals who are symptomatic. Interestingly, asymptomatic mutation-carriers do not display these same characteristics at their baseline visits. By following these individuals as they progress towards their EYO, the DIAN Observational Study offers an opportunity for researchers to track their inevitable structural decline in a manner that has not previously been possible. Given that reduction in grey matter volume and thickness, increased Aβ deposition, and disturbed glucose metabolism are all hallmark characteristics of AD-related neuropathology, these observations of the DIAN cohort are clear evidence that this study can make meaningful contributions to our understanding of both ADAD and sporadic AD.

Beyond neuroimaging, the DIAN Observational Study also involves seven major cores, who work together to ensure the collection of a large number of additional data modes (Figure 10). Almost all the participants outlined in the current paper have also had extensive genetic, clinical, cognitive, and fluid biomarker samples collected. Genetic data consists of mutations status in the *APP*, *PSEN1* and *PSEN2* genes, as well as information about AD-associated single nucleotide polymorphisms (SNPs) such as the well-studied variants in the genes coding for the *APOE* ^140^ and brain-derived neurotrophic factor (*BDNF*) proteins ^95, 141^. Whole genome sequencing of individuals also allows for the possibility of conducting much more extensive exploratory genetic analyses on individuals within this cohort ^142^. In addition to genetic information, participants also agreed to provide biological samples such as CSF and blood plasma. Using these measures, it is of particular interest to quantify levels of fluid biomarkers such as the 40 and 42 amino acid forms of the Aβ peptide (Aβ40 and Aβ42, respectively), which can be used to infer ratios of Aβ forms within the brain ^143^. Similarly, phosphorylated tau181 (pTau) and total tau (tTau) measured from these samples are indirect measures of amyloid plaques ^144^ that precede the appearance of the intraneuronal neurofibrillary tangles comprised primarily of tau ^145, 146^. Beyond these traditional markers of Aβ and tau, other fluid biomarkers of interest such as CSF- and plasma-derived concentrations of the neurofilament light (NfL) and soluble triggering receptor expressed on myeloid cells 2 (sTREM2) proteins, have recently been added to the DIAN Observational Study as they have been identified as having robust diagnostic and prognostic utility for understanding AD trajectory ^145, 147, 148^. Additionally, DIAN participants also complete a large number of cognitive tasks, in order to have their cognitive abilities extensively phenotyped. Many of the tasks administered are common neuropsychological batteries such as the revised Wechsler Adult Intelligence Scale (WAIS-R; ^149^) and Wechsler Memory Scale (WMS; ^150^), which aim to characterize memory, attention, processing speed, and inhibition. These are critical components of cognition previously reported to decline in association with AD ^151^.

**Figure 10.**
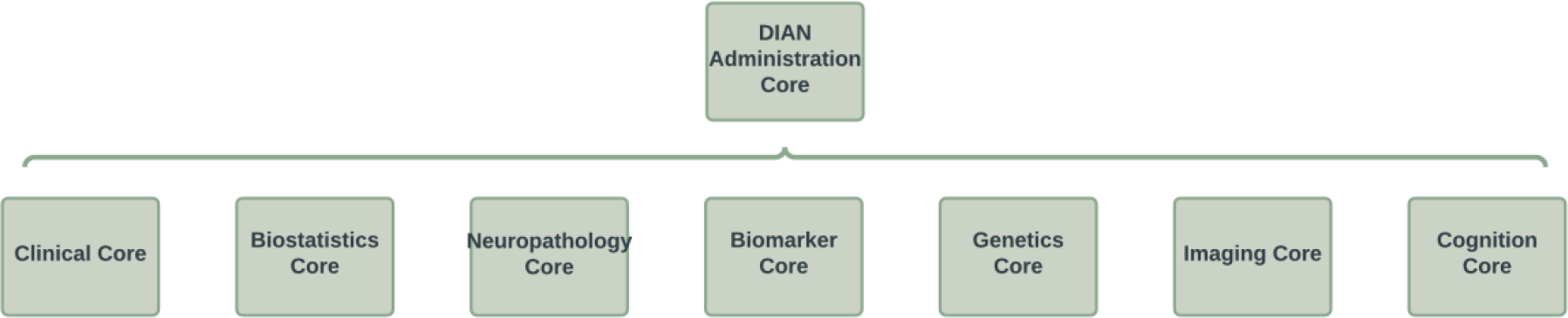
Structural outline of the collaborating cores of the DIAN Observational Study. The DIAN study is comprised of several core centers, each working symbiotically to collect, analyze, and prepare data for regular data freeze releases.

Finally, some DIAN participants have also opted to donate their brain tissues upon death. This extraordinary donation (a valuable resource that will increase with time) allows for fine grained investigation of the neuropathology of end-stage AD, facilitates important post-mortem validation of neuroimaging and biofluid studies done *in vivo,* and enables cutting-edge molecular analyses such as single-nuclei RNA seq, proteomics, lipidomics, et cetera.

Similar to the neuroimaging data described previously, much of these cross-modal data are available by request. These additional data position the DIAN Observational Study uniquely as a deeply phenotyped cohort, with many opportunities to advance the understanding of the trajectory of preclinical to diagnosed AD. Beyond ADAD, the DIAN dataset also provides a unique set of deeply-phenotyped middle-aged healthy controls. This cohort of individuals is important given the relative scarcity of data available for non-clinical individuals of this age range, an issue that has historically been noted in studies aiming to understand brain changes across the lifespan.

To date, 175 studies have been published using DIAN data. This highlights the importance of the DIAN Observational Study for shaping our knowledge of preclinical AD, and by extension, sporadic AD. A major outcome of the DIAN Observational Study has been its use in furthering our understanding of the temporal order of biomarker changes that occur in the lead up to clinically diagnosed AD. Bateman et al. (2012) determined that ADAD was associated with a series of pathological changes in brain structure and function that begin as early as 20 years prior to typical AD diagnosis. While this seminal work was conducted using data from ADAD individuals, it is thought these same patterns of biomarker changes could be present in the lead up to the development of sporadic AD. Using the DIAN Observational Study as a “model” for understanding sporadic AD allows us to get a clearer framework for what is going on within the system prior to overt cognitive decline, which can often be the first sign of disease in those with sporadic AD. The neuroimaging data collected via the DIAN Observational Study have also informed several other key areas of AD research including better understanding the similarities and differences in grey matter atrophy for the sporadic and autosomal dominant forms of AD ^33, 65^. These studies have highlighted that regions such as the hippocampus, traditionally thought to be highly predictive of AD, index non-specific brain atrophy ^152^. In contrast, cortical thickness has been reported to be a sensitive measure of AD-related grey matter pathology ^137, 139^. Building upon this idea, Dincer et al. (2020) used DIAN data to propose a cortical signature of AD-related atrophy, highlighting that while there are differences in the cortical thinning patterns of ADAD and sporadic AD, the spatial patterns of AD-related volumetric decline in these two cohorts share many similarities. In addition to expanding our understanding of established biomarkers, other work highlighting data from the DIAN cohort have focused on validating emerging biomarkers of AD-related pathology ^143, 145, 148^. For example, work has recently reported that increases in NfL protein concentrations are correlated with decreases in white matter integrity ^148^, with rate of change in the correlation of this protein sensitive enough to discriminate mutation carriers from non-carriers ^145^. The DIAN Observational Study has also made it possible to study a number of other biomarkers such as CSF neurogranin, synaptosomal-associated protein-25 (SNAP-25), visinin-like protein 1 (VLIP-1), and chitinase-3-like protein 1 (YKL-40). These proteins are known to be associated with synaptic damage, neuronal injury, and neuroinflammation, all of which accrue following the onset of amyloid deposition ^143^.

Importantly, the DIAN Observational Study has been critical for providing the scientific framework that allowed for the formation of the DIAN-TU. Beginning in 2012, the DIAN-TU was one of the world’s first collaborations focusing on the prevention of dominantly inherited AD. To this end, the DIAN-TU enrolls individuals from families who carry genetic mutations that lead to ADAD, many of whom may have participated in the DIAN Observational Study. The main goal of the DIAN-TU is to assess the safety, tolerability, and effectiveness of drugs that may improve the lives of those at-risk or living with AD. These studies will help researchers understand whether these drugs can be used to prevent, delay, or even reverse the neuropathological changes that occur in dominantly inherited AD. To date, the DIAN-TU has enrolled 228 individuals from across the globe and is actively running trials to assess drugs targeting amyloid and tau deposition within the brain. The DIAN Observational Study was fundamental for the inception of this trial, as it allowed for researchers to demonstrate the statistically powerful design inherent in this cohort.

Finally, a major goal of the DIAN Observational Study was to create a large data resource to be shared across AD researchers around the globe. To that end, the data outlined in this manuscript are available by request at https://dian.wustl.edu/our-research/observational-study/dian-observational-study-investigator-resources/. Importantly, all available data are organized in a way that preserves participant privacy. At the point of writing, DIAN Data Freeze 15 is available; and as the DIAN Observational Study continues, additional data will be released biannually. Major efforts are ongoing around the world to attempt to understand AD, in order to inform diagnoses, treatments, and potential cures.

Given the rich dataset that the DIAN Observational Study is generating, the authors hope this resource manuscript will help to outline the neuroimaging data that can be requested from the DIAN Observational Study.

## Acknowledgements

Data collection and sharing for this project was supported by the Dominantly Inherited Alzheimer Network (DIAN, U19AG032438) funded by the National Institute on Aging (NIA), the Alzheimer’s Association (SG-20-690363-DIAN), the German Center for Neurodegenerative Diseases (DZNE), Raul Carrea Institute for Neurological Research (FLENI). Partial support by the Research and Development Grants for Dementia from Japan Agency for Medical Research and Development, AMED, and the Korea Health Technology R&D Project through Korea Health Industry Development Institute (KHIDI), Spanish Institute of Health Carlos III (ISCIII), Canadian Institutes of Health Research (CIHR), Canadian Consortium of Neurodegeneration and Aging, Brain Canada Foundation and Fonds de Recherche du Québec – Santé. This manuscript has been reviewed by DIAN Study investigators for scientific content and consistency of data interpretation with previous DIAN Study publications. We acknowledge the altruism of the participants and their families, and contributions of the DIAN research and support staff at each of the participating sites for their contributions to this study.

## Data Availability Statement

The datasets analyzed during the current study are available following the completion of a Dominantly Inherited Alzheimer Network Data Request, [https://dian.wustl.edu/our-research/for-investigators/dian-observational-study-investigator-resources/data-request-terms-and-instructions/].

## Code Availability Statement

Upon publication, the analysis scripts used to generate the images and statistical output in this manuscript are available from the Benzinger Gitlab Repository, [https://github.com/benzinger-icl]. These scripts use openly accessible packages within the R (version 4.1.2) environment.

## Ethics Statement

All study procedures were approved by the Washington University Human Research Protection Office and the local institutional review boards of the participating sites. Furthermore, all participants recruited into the DIAN study provided informed consent.

